# Telomere variant sequences encode the genetic blueprint for allele-specific telomere length

**DOI:** 10.64898/2026.08.04.741677

**Authors:** Xiaoran Chai, LaiFong Poon, Hengrui Liu, Manvendra K. Singh, Takashi Minami, Goro Sashida, Masafumi Fukuda, Bin Tean Teh, Patrick Tan, William Ying Khee Hwang, Jue Lin, Angela S. Koh, Tatsuro Kondoh, Lifeng Xu, Motomi Osato, Jin Liu, Shang Li

## Abstract

The ends of human chromosomes are capped by specialized nucleoprotein structures known as telomeres, which are essential for genome stability. Recent advances in long-read sequencing have enabled allele-specific telomere length measurements at nucleotide resolution, uncovering extreme heterogeneity in telomere length between alleles that is determined at birth and gradually shortens with age. However, the mechanisms underlying allele-specific telomere maintenance and its stability remain poorly understood. Here, we developed a high-resolution workflow combining PacBio and Nanopore long-read sequencing platforms to map allele-specific telomere length in clinical patient samples as well as telomerase-positive cancer cell lines. By tracing allele-specific telomeric sequence in family members across multiple generations, we show that telomeric variant sequences (TVSs) interspersed throughout the canonical repeat region are heritable (with mean similarity score > 0.95), allele-specific, and account for the extreme heterogeneity of telomere length between alleles. Targeted deletion of allele-specific TVSs using CRISPR-Cas9 resets telomere length, further confirming their causal role in the control of allele-specific telomere maintenance. Continuous cell proliferation likely drives the slow but stochastic evolution of allele-specific TVSs distribution, resulting in asymmetry in telomere inheritance from father and mother (p-value < 0.045). These results illustrate how the telomere inheritance may predict allele-specific vulnerability in telomere protection, providing a new paradigm for personalized telomere profiling in aging-related diseases.

## Introduction

The ends of human chromosomes are capped by specialized nucleoprotein structures called telomeres, which protect chromosome termini from degradation, end-to-end fusion, and recombination (*1, 2*). Telomeres consist of long tracts of double-stranded hexa-nucleotide DNA repeats (5’-TTAGGG-3’)n, terminating in single-stranded 3’ G-rich overhangs (*3, 4*). These overhangs can fold back to form a protective telomeric loop (T-loop) (*5*).

Telomeric DNA repeats are bound by a six-protein complex known as shelterin, which distinguishes natural chromosomal ends from DNA double-strand breaks. The core shelterin components include TRF1, TRF2, TIN2, POT1, TPP1, and RAP1 (*6–11*). TRF1 and TRF2 bind the double-stranded canonical telomere motif 5’-TTAGGGTTA-3’, whereas POT1 binds the 3’ single-stranded overhang (*6–11*). Even a single nucleotide substitution within the canonical sequence can drastically reduce TRF1 and TRF2 binding affinity (*11, 12*), compromising end protection and triggering telomere dysfunction-induced foci (TIFs) (*11–13*). In addition to safeguarding telomeres, shelterin plays an essential role in regulating telomere length homeostasis. Telomere-binding proteins on longer telomeres may promote erosion, inhibit elongation, or both (*14–16*), thereby progressively limiting telomerase access. As a result, telomere length in telomerase-positive cells is maintained by a dynamic balance between telomerase-mediated extension and shelterin-mediated suppression. This “protein counting” model is well supported across species, with TRF1, TRF2, and POT1 functioning as negative regulators of telomere length (*15, 17, 18*).

Telomeric DNA is synthesized by the enzyme telomerase, which comprises two essential subunits: the protein component hTERT and the RNA component hTR (*19, 20*). While hTR is ubiquitously expressed, *hTERT* expression is the rate-limiting factor and is barely detectable in most adult somatic tissues, except in germ cells and tissue stem cells (*19–23*). In somatic cells lacking telomerase activity, telomeres progressively shorten by approximately 50–200 bp per cell division due to incomplete replication of chromosome ends and end-processing events (*4, 24*). When telomeres become critically short, they elicit DNA damage responses (*11–13*). Just a few of these short telomeres are sufficient to trigger senescence or apoptosis in human cells that is associated with human aging-related degenerative defects (*25, 26*). Thus, telomere length at birth and the cumulative telomere attrition across the lifespan limit the replicative potential of somatic cells during aging (*27, 28*).

Although telomere length is maintained in pluripotent stem cells, telomerase activity in tissue-specific progenitor and stem cells is generally insufficient for long-term telomere maintenance (*29*). Consequently, telomere shortening also occurs in these compartments, progressively limiting their proliferative capacity and contributing directly to age-associated functional decline (*29–31*). Moreover, genetic mutations in telomere-and telomerase-associated genes underlie a spectrum of inherited disorders, collectively known as telomere syndromes or telomeropathies (*32, 33*), which are characterized by reduced telomerase activity, accelerated telomere shortening, and premature aging phenotypes. These observations underscore the importance of telomere homeostasis in human health and highlight the potential of telomere length as a predictive biomarker for aging and disease risk.

Allele-specific telomere length distributions have been observed in individuals, defined in the zygote and maintained throughout life (*34, 35*). Long-read sequencing has recently revealed extreme heterogeneity in allele-specific telomere length (*36–38*), apparently established at birth and gradually shortening with age (*36*). Yet, the molecular mechanisms underlying this allelic disparity in telomere maintenance remain poorly understood. Using our recently established long-read sequencing approach for high-throughput telomere profiling (*39*), we also identified distinctive structural features, most notably telomeric variant sequences (TVSs) interspersed throughout the telomeric repeat tracts. Each chromosomal end carries a unique pattern of TVSs, with larger and more abundant variants typically located near the subtelomeric boundary. These unique patterns are individual-specific and not associated with ethnicity. Because TVSs disrupt the canonical telomeric repeat sequence, they can reduce shelterin binding density and contribute to variation in telomere protection during aging (*39*). While prior studies using long-read whole-genome data have hinted that telomeric variant repeats from specific chromosome ends may be inherited (*40*), low sequencing depth has precluded confirmation of this for all the chromosome ends. Furthermore, the stability and heritability of allele-specific TVS distributions have not been fully established.

Here, we have combined PacBio and Nanopore long-read sequencing platforms to map allele-specific telomere length in patient samples and telomerase-positive cancer cell lines. We demonstrated that TVSs dispersed along the telomere repeat region are heritable and allele-specific, and account for the extreme heterogeneity of telomere length between alleles. The presence of allele-specific telomere lengths and TVS distributions further suggests that the propensity for telomere uncapping—and thus cellular aging or genome instability—may vary between alleles depending on their specific TVS architecture.

## Results

### Telomere-enriched long-read sequencing enables precise allele-specific telomere mapping

Our previous studies revealed extensive heterogeneity in TVSs, which include both telomere variant repeats (TVRs) and non-repeat sequences ranging from one to several hundred nucleotides (*39*). These variants are interspersed throughout the telomeric repeat region, with larger and more abundant TVSs typically located near the subtelomeric end (**Figure 1A**). While the allele-specific distribution and sequence of TVSs are unique to each individual, the stability and heritability of these patterns across generations have not yet been established. Therefore, it remains unclear how inherited allele-specific TVSs distribution influences the inherited telomere maintenance, predicts allele-specific vulnerability in telomere protection, and acts in aging-related diseases.

**Figure 1:**
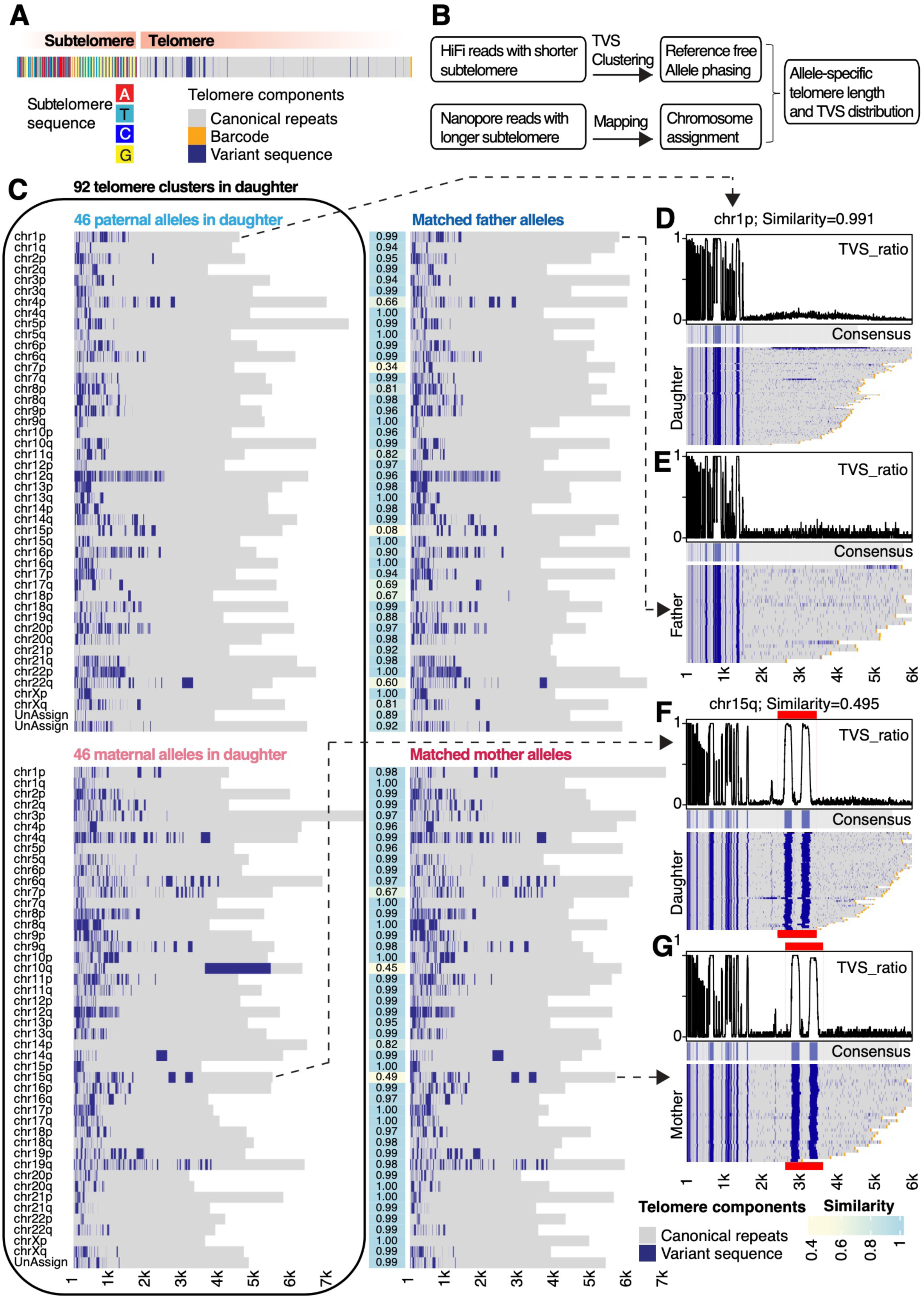
Allele-specific telomere variant sequences (TVSs) are stable and inheritable traits in the offspring of family 1. Allele-specific TVS patterns in a trio from Family 1: father, mother, and daughter. **A)** A telomere-containing read can be divided into two parts: the telomeric region and the subtelomeric region. The telomere start site is defined as the first occurrence of “TTAGGGTTAGGG”. The telomeric region is primarily composed of canonical telomere repeats (gray), interspersed with telomere variant sequences (TVSs, dark blue). **B)** Schematic overview of the analysis pipeline, including reference-free allele phasing of telomere-enriched PacBio HiFi reads, and chromosome assignment of Nanopore/PacBio Revio whole-genome sequencing (WGS) reads by mapping onto the reference genomes. **C)** TVS profiles of 92 allele-specific telomere clusters identified in the daughter’s peripheral blood leukocytes (PBLs), comprising 46 paternally inherited and 46 maternally inherited alleles (left), and the corresponding TVS-matched alleles from the father and mother (right). Pairwise TVS similarities (Pearson correlation of TVS ratios) are visualized as a heatmap (middle). **D**–**E)** Allele-specific reads from Chr. 1p in the daughter (81 reads, D) and father (26 reads, E). In each group, the TVS ratio at each position is plotted as a line graph to visualize allele-specific TVS profiles. **F**–**G)** Allele-specific reads from Chr. 15q in the daughter (63 reads, F) and mother (32 reads, G). Shifts in TVS blocks (red segments) are shown.

To address this knowledge gap, we developed a high-resolution workflow combining PacBio and Nanopore long-read sequencing platforms to map allele-specific telomere length in clinical patient samples as well as telomerase-positive cancer cell lines. To enable both accurate telomere length measurement and robust allele-specific mapping, two distinct DNA libraries were prepared for each sample. The first library, enriched for telomeric DNA via Telobait and endonuclease digestion with a combination of RsaI, HinfI, and EcoRI, was sequenced using PacBio HiFi for accurate telomere profiling. However, enzymatic digestion may truncate subtelomeric regions (sometimes to < 50 bp) compromising chromosomal end assignment. To overcome this issue, a second, unenriched WGS library was sequenced using PacBio Revio and/or Nanopore to provide longer reads that capture full subtelomeric regions and allele-specific TVSs distributions at lower read depth. This dual-sequencing approach (**Figure 1B**) significantly improved allele-specific mapping accuracy, and enabled reference-free phasing of telomere-containing reads (**Supplementary Figure S1**). HiFi-enriched telomere reads were partitioned into allele-specific groups based on their unique TVS distributions using a two-step process: dimensionality reduction via t-SNE (*41*), followed by noise-aware clustering with HDBSCAN on the top two t-SNE components (*42*) (see Methods). In parallel, WGS-derived telomere reads were also grouped into phased alleles based on their TVS patterns.

Next, we mapped the WGS reads to a customized reference genome combining T2T-CHM13v2.0 (*43*) and the recently released human pangenome (*44*). While preserving robust allele phasing of the telomere-enriched data, chromosome assignments from WGS reads can be transferred to the corresponding telomere-enriched HiFi reads. The power of this approach is demonstrated in the Chr. 10p analysis (**Supplementary Figure S1**), where the two telomeric alleles (distinguished by distinct TVS patterns) were unambiguously clustered into separate haplotypes. By aligning reads from both sequencing libraries (telomere-enriched HiFi sequencing and WGS) using the telomere start site (first “TTAGGGTTAGGG”) as a common anchor, we could confidently match the phased HiFi clusters to their corresponding mapped WGS alleles. Overall, this pipeline yielded high-confidence allele-resolved maps of telomeric sequence and structure, suitable for downstream quantitative analysis.

### Allele-specific TVSs are stable and inheritable traits

To investigate the stability and heritability of TVSs across generations, we collected peripheral blood leukocytes (PBLs) from three independent families and performed telomere long-read sequencing. For Family 1, 92, 90, and 90 allele-specific telomere clusters were identified in the daughter, father, and mother, respectively (**Figure 1C; Supplementary Figures S2-S3; and Extended Data 1**). Most clusters could be confidently assigned to specific chromosomal ends. Nevertheless, and despite the overall robustness of this strategy, some telomeric alleles remained unassigned or unresolved, likely due to internal restriction enzyme cutting within the telomeric region that eliminates the subtelomeric anchor needed for allele mapping, or complex repeat architecture in certain subtelomeric regions (**Supplementary Figures S2–S5**).

Although telomere-containing reads from the same allele may exhibit minor variation due to stochastic DNA replication errors, their TVS patterns remained highly similar (**Figure 1C; Supplementary Figures S2-S3**). To quantify these distributions, we calculated a positional TVS ratio for each allele, defined as the fraction of reads at a given position that contain a TVS. We then plotted this ratio across the length of the telomere to generate a unique TVS signature (**Figure 1D–1G; and Extended Data 1**). Then, we constructed a consensus TVS sequence for each allele using a majority rule: positions with a TVS ratio >0.5 were designated as variant, and others as canonical. This framework enabled quantitative comparison of TVS patterns across generations. We computed a similarity score between alleles based on the correlation of their TVS ratios starting from the telomere start site. Using this metric, we identified 46 paternally and maternally inherited alleles in the daughter from Family 1. Most of these displayed high similarity to the corresponding parental alleles (**Figure 1C–1E**). For example, the daughter’s Chr. 1p allele was nearly identical to the paternal allele, with a similarity score of 0.99.

In some cases, we noted a divergence in inherited TVS patterns. For example, the daughter’s Chr. 15p allele, while similar to the maternal version, showed altered TVS spacing, yielding a low similarity score of 0.495 (**Figure 1F–1G**). Likewise, the daughter’s Chr. 10q allele exhibited a new >1 kb stretch of TVSs composed predominantly of (TCAGGG)n repeats, which was not present in the mother’s corresponding allele (**Figure 1C; and Extended Data 1**). Notably, this sequence patch was only present in a subset of telomere-containing reads, suggesting a recent somatic event in hematopoietic stem cells. Overall, these results indicate that allele-specific TVS patterns are highly stable and heritable across one generation. We posit that occasional deviations likely result from stochastic mutation or recombination events during DNA replication.

To extend our findings across multiple generations, we analyzed Family 2, comprising a grandfather, grandmother, mother, and daughter. Using our allele-resolved telomere sequencing pipeline, we identified 88 telomeric alleles in the mother, of which 45 were traced to the grandfather and 43 to the grandmother. In the daughter, we identified 44 maternally inherited alleles, 28 originating from the grandfather and 16 from the grandmother (**Figure 2A-2D; Extended Data 2; and Supplementary Figure S4**). Similar results were obtained in Family 3, confirming the stability and heritability of TVS distributions across lineages (**Supplementary Figure S5**).

**Figure 2:**
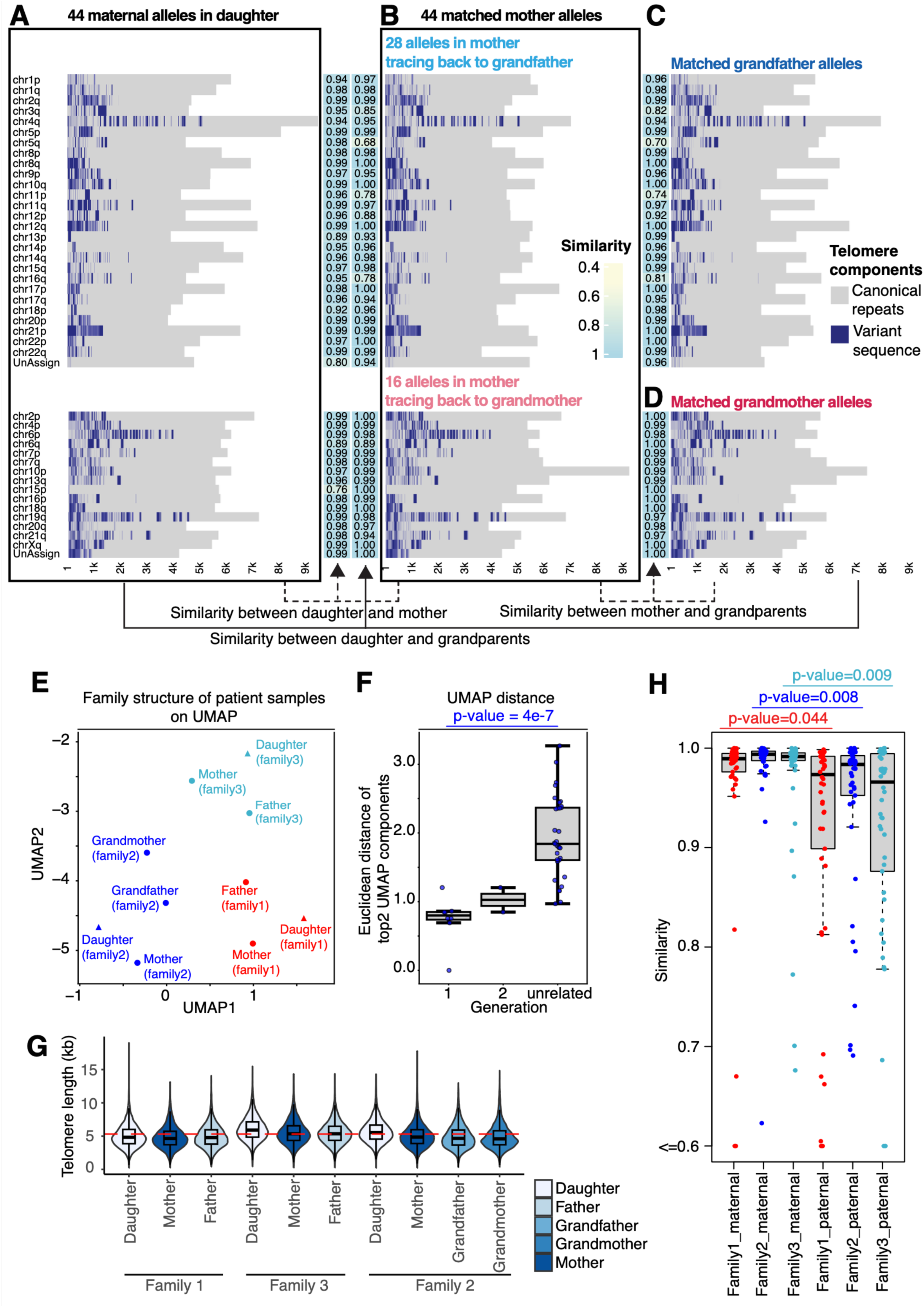
Allele-specific telomere variant sequences (TVSs) are stably inherited across multiple generations. **A**–**D)** Allele-specific TVS patterns in a three-generation family (Family 2) comprising daughter, mother, grandfather, and grandmother. **A)** TVS distributions in 44 maternal alleles identified in the daughter’s peripheral blood leukocytes. **B)** The 44 maternally inherited alleles in the daughter correspond to 44 alleles in the mother, 28 of which can be traced to the grandfather (**C)** and 16 to the grandmother (**D)**. Heatmaps between panels show pairwise TVS similarity scores for each inherited allele. **E)** UMAP projection illustrating family structure across all individuals based on global TVS similarity. **F)** The Euclidean distance between samples in the first two UMAP dimensions and the correlation with generational proximity. **G)** Telomere length distributions in all individuals across the three families. The dotted red line indicates the mean telomere length across all the samples. **H)** Comparison of similarity scores between paternally and maternally inherited alleles across three independent families. **F)**

We further visualized the relatedness of TVS patterns using UMAP dimensionality reduction. TVS profiles clustered by family, with intra-generational similarity reflected in closer distances between samples (**Figure 2E**). Parent–offspring pairs showed the smallest Euclidean distances, followed by grandparent–grandchild pairs, while unrelated individuals were the most divergent (**Figure 2F**). In line with known biology, mean telomere length shows an age-dependent decline (**Figure 2G**).

Our high-resolution data also captured instances of chromosomal crossover and mitotic recombination between telomeric regions from different chromosomal ends. For example, Chr. 17q in the mother from Family 2 contained a TVS patch matching the grandfather’s allele—despite not being inherited by the mother as a whole (**Supplementary Figures S4 and S6**). Similarly, the mother’s Chr. 11p from Family2 contained a TVS patch resembling the grandmother’s Chr. 12p allele (**Supplementary Figures S4 and S7**). These events are likely the outcome of the meiotic or mitotic recombination (**Supplementary Figure S8**). Overall, our method not only elucidates the stable transmission of TVSs, but also identifies rare recombination events at telomeres during inheritance.

### Parental origin affects the stability of inherited TVS patterns

Having established the overall stability of TVS patterns across generations, we next investigated whether this stability differed based on parental origin. In Family 1, direct comparison of similarity scores revealed a significant parent-of-origin effect: paternally inherited alleles were significantly less stable than maternally inherited ones (p = 0.044; **Figure 2H**). Notably, 13 of 46 paternally inherited alleles showed similarity scores < 0.9, compared to only 4 of 46 maternally inherited alleles. A similar trend was observed in Family 2 during the first generational transmission (grandparents to mother), where paternally inherited alleles again exhibited significantly lower stability (p = 0.008; **Figure 2H**) than maternally inherited ones. This multi-generational comparison confirmed the parent-of-origin effect with high statistical power. Subsequent transmission from mother to daughter exhibited very high stability, with only 4 of 44 inherited alleles falling below the 0.9 similarity threshold (**Figure 2A–2B**). We consistently obtained similar results from Family 3 (p = 0.009; **Figure 2H and Supplementary Figure S5**).

These results confirm both the general stability of TVS signatures across generations and a clear asymmetry in telomere inheritance. Paternal alleles are more prone to divergence, likely due to the continuous nature of spermatogenesis and the accumulation of replication-associated errors in male germ cells. By contrast, maternal oocytes remain arrested in meiosis I until ovulation, preserving TVS integrity. This mechanistic difference may underlie the observed bias in TVS stability by parental origin.

### Allele-specific TVSs underlie the allele-specific telomere length

The allele-specific telomere length data from PBLs in the three families highlighted a potential positive correlation between telomere length and TVS abundance. In primary lymphocytes, however, these correlations were relatively modest (r = 0.55, r = 0.70, and r = 0.56 for the daughter, father, and mother in Family 1, respectively; **Figures 3A–3C**), and similar patterns were seen in Families 2 and 3 (**Supplementary Figure S9**). This finding likely reflects the stochastic nature of telomere shortening in hematopoietic stem and progenitor cells, which have low telomerase activity and therefore accumulate heterogeneity in telomere length over time. For example, in Family 1, the daughter inherited a Chr. 5p telomere from the father that is almost 4 kb longer than the Chr. 20p telomere inherited from the mother (**Figure 1C**). By contrast, telomere length is maintained in telomerase-positive cancer cell lines via a homeostatic balance between telomerase-mediated extension and shelterin-mediated inhibition. This led us to hypothesize that the influence of TVSs on allele-specific telomere length would be more evident in telomerase-positive cancer cells.

**Figure 3:**
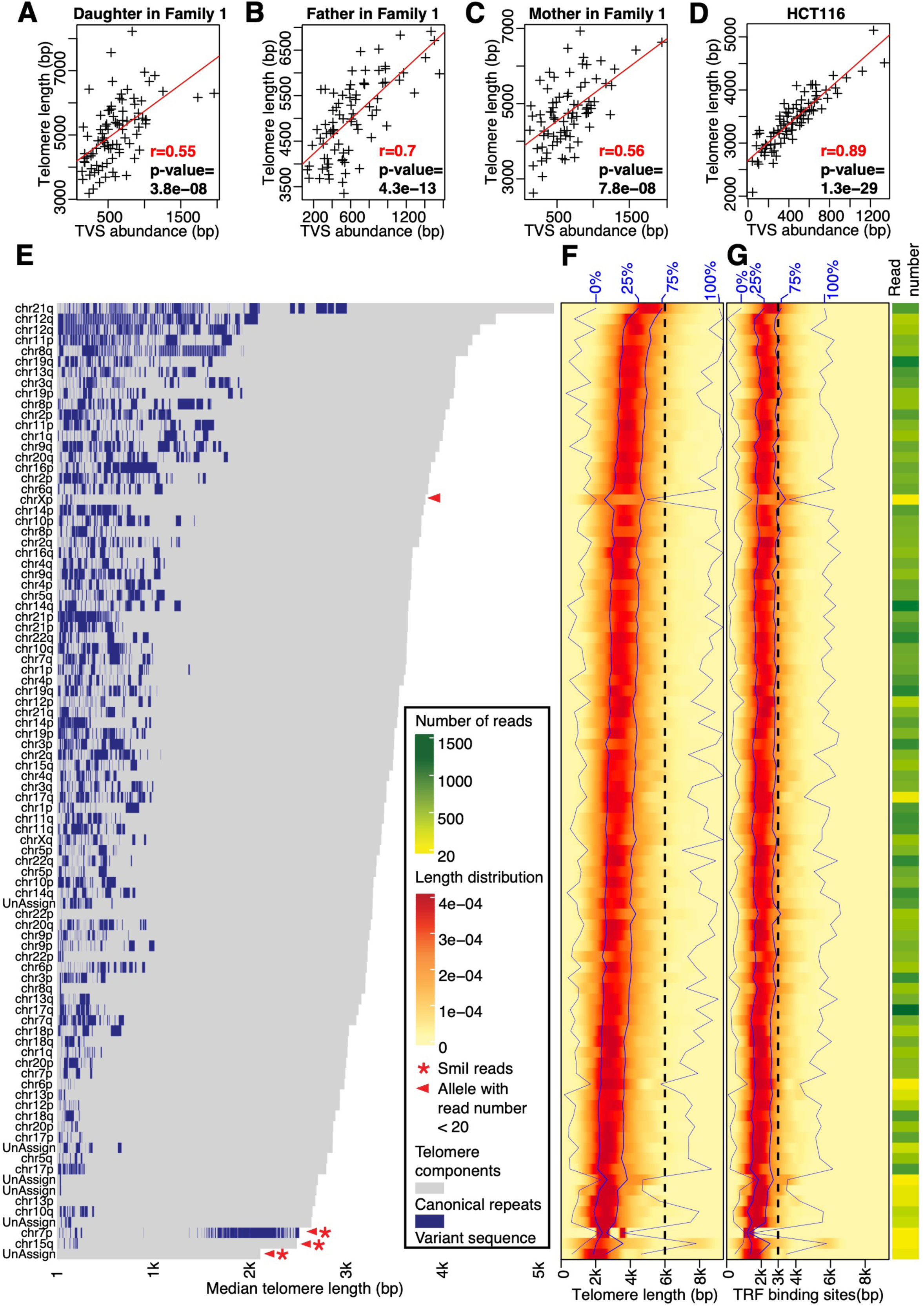
Allele-specific telomere length is positively correlated with TVSs abundance in HCT116. **A**–**D)** Correlation between allele-specific telomere length and TVSs abundance in peripheral blood leukocytes (PBLs) of Family 1 (daughter, father, and mother) and in the telomerase-positive cancer cell line HCT116. **E)** Allele-specific telomere lengths in HCT116, in order from longest to shortest. **F)** The heterogeneity of allele-specific telomere length in HCT116 cells. **G)** The distributions of the maximum number of potential TRF1/TRF2 binding motifs (TTAGGGTTA repeats) per allele plotted alongside telomere lengths (shown in F). Most allele clusters were generated using the RsaI/HinfI/EcoRI enzyme combination during the telomere enrichment step. However, three clusters (red stars) were generated using BseJI/SmiI, Eco32I/SmiI or StuI/SmiI. Clusters with <20 reads (red triangles) were excluded from correlation analyses between telomere length and TVS abundance.

Supporting this, analysis of the HCT116 cell line revealed a strong positive correlation between TVS abundance and allele-specific telomere length (r = 0.89, p = 4.2e-30; **Figure 3D**). Across alleles, telomere lengths spanned more than 3 kb in HCT116 cells, and the correlation remained visually and statistically apparent when alleles were ranked by mean length (**Figures 3E-3F**). These findings suggest that TVS abundance plays a key role in modulating allele-specific telomere length in telomerase-positive cells.

Given the known role of telomere “protein counting” mechanisms, binding of shelterin complexes regulate telomerase access (*14–16*), we further assessed the relationship between TRF1/TRF2 motif content and telomere length in HCT116 cells. TRF binding sites were quantified as non-overlapping “TTAGGGTTA” motifs within the telomeric region. Although TVSs may interrupt the protein binding of the sheltering complex by disrupting the continuity of the telomere canonical repeats, the number of TRF1/TRF2 binding sites per allele remained relatively constant across HCT116 alleles (**Figure 3G**). Specifically, the standard deviation (SD) of TRF site counts was 13 (equivalent to ∼117 bp), whereas the SD of telomere length was 495 bp. Similarly, the coefficient of variation (CV) for TRF site counts was 8.2%, nearly half that of telomere length (14.4%). Collectively, the number of canonical protein-binding motifs is not significantly altered, despite the introduction of variant sequences. These data suggest that while TVSs define telomere length heterogeneity, TRF1/TRF2 binding site density remains buffered, potentially stabilizing shelterin-mediated telomere length regulation.

To further explore the dynamics of TVS-dependent telomere regulation, we profiled allele-specific telomere length in three single-cell derived clones of the HT1080 cell line, each exhibiting distinct mean telomere length (**Figure 4A-4D, and Supplementary Figure S10**). This approach allowed us to track the impact of TVS distribution during the reset (establishment of new telomere length baselines) of allele-specific telomere length in each progeny from parental HT1080. Clones A6 and B3 had similar mean telomere lengths, while clone B2 displayed markedly shorter mean telomeres by ∼4kb. Despite this difference, the allele-specific telomere lengths across clones preserved their relative ranking (i.e., long vs. short alleles remained consistent), and this ranking closely reflected the overall mean telomere length of each clone (**Figure 4A-4D, and Supplementary Figure S10A and S10C**). By contrast, while absolute telomere lengths varied across clones, the TVS patterns for any given allele remained stable and nearly identical between A6, B3, and B2 (**Figure 4A-4C, and Supplementary Figure S10A-S10B**). Importantly, TVS abundance strongly predicted telomere length within each clone, particularly in clone B2 (r=0.81, p=5.3e-23; **Figure 4E-4G**), which has shortest telomere compared to A6 and B3, further supporting the hypothesis that TVS content governs telomere length dynamics in telomerase-positive cells. Notably, in clone B2, we observed the loss of 3’ TVS blocks in eight alleles (likely due to telomere re-setting during clonal expansion), and only a single case of TVS gain (**Supplementary Figure S10D-10F**).

**Figure 4:**
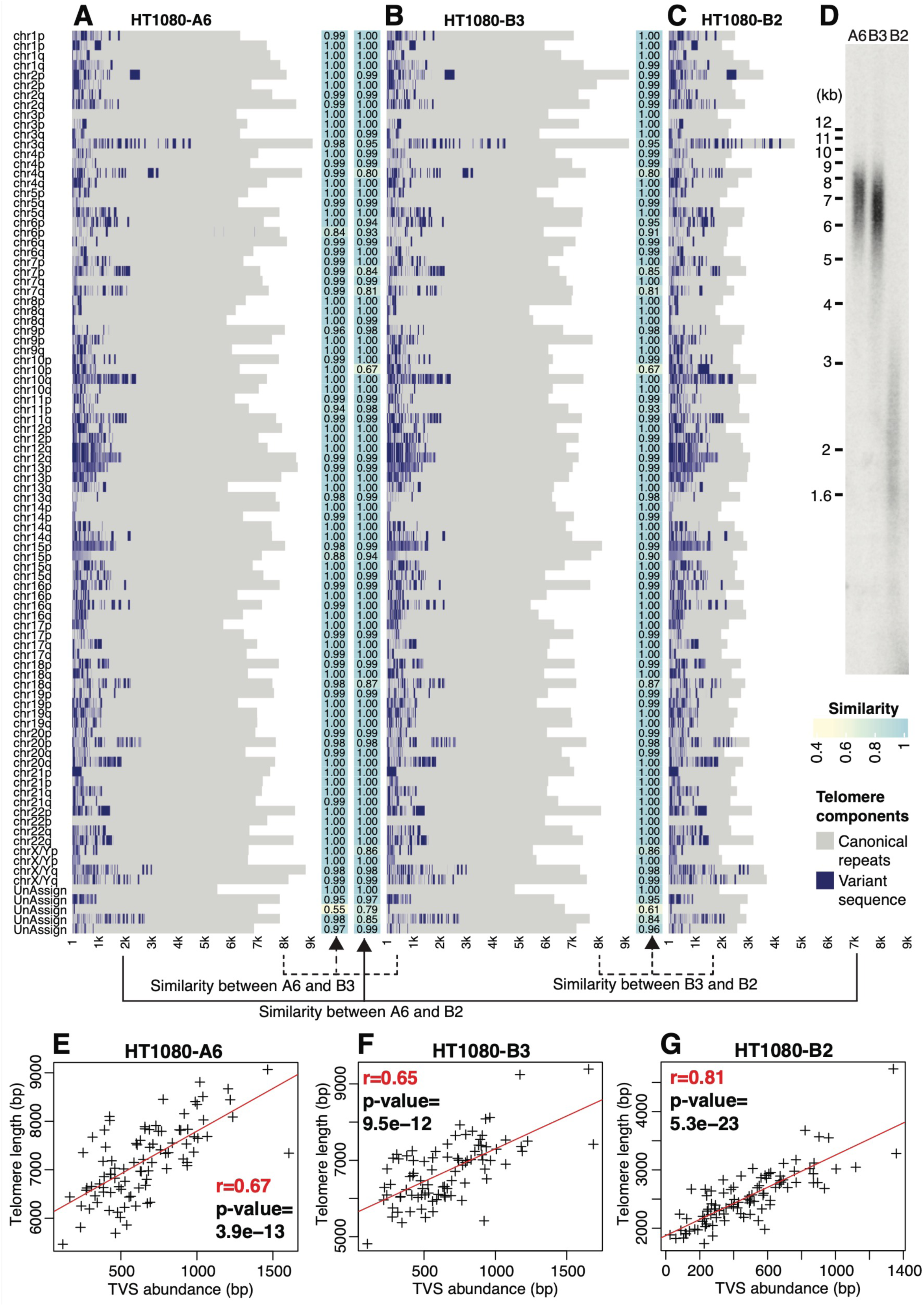
TVS abundance correlates with allele-specific telomere length in single-cell-derived HT1080 subclones with distinct mean telomere lengths. **A–C)** Allele-specific telomere lengths across all 92 alleles in three single-cell-derived HT1080 subclones: HT1080-A6, HT1080-B3, and HT1080-B2. **D)** Teloblot analysis showing the overall telomere length distribution in the three HT1080 subclones. **E–G)** Correlation between allele-specific telomere length and TVS abundance in the three single-cell derived HT1080 subclones.

Together, these results suggest that in telomerase-positive cancer cells, TVS abundance is an intrinsic determinant of allele-specific telomere length by modulating TRF1/TRF2 binding site density. This model reconciles the observed variability in telomere length with the relative stability of shelterin-mediated counting across telomeres.

### Allele-specific telomere length can be reset by adjusting TVSs abundance

To directly test whether the TVS distribution dictates allele-specific telomere length, we used CRISPR-Cas9 to delete TVS patches in a selected allele of HCT116 cells. We designed sgRNA targeting unique TVS blocks on the Chr. 1q allele 1 (Chr. 1q-A1; **Figure 5A**). Following CRISPR editing, single-cell-derived clones were isolated and cultured for > 50 population doublings to allow telomere lengths to reach a new steady state as the telomerase activity in HCT116 can restore canonical terminal repeats after TVS deletion (**Figure 5C, and Supplementary Figures S11A–S11B**). Long-read sequencing confirmed the cutting of the TVSs in Chr. 1q-A1 at both sgRNA sites (Target 1 and Target 2, **Figure 5A**). Deletion at Target 1 resulted in the loss of nearly all TVSs in Chr. 1q-A1 (clone 2 and 3, **Figure 5A-5C**), whereas editing at Target 2 had minimal effect on TVS abundance (clone 1, **Figure 5A; and Supplementary Figure S11C**).

**Figure 5:**
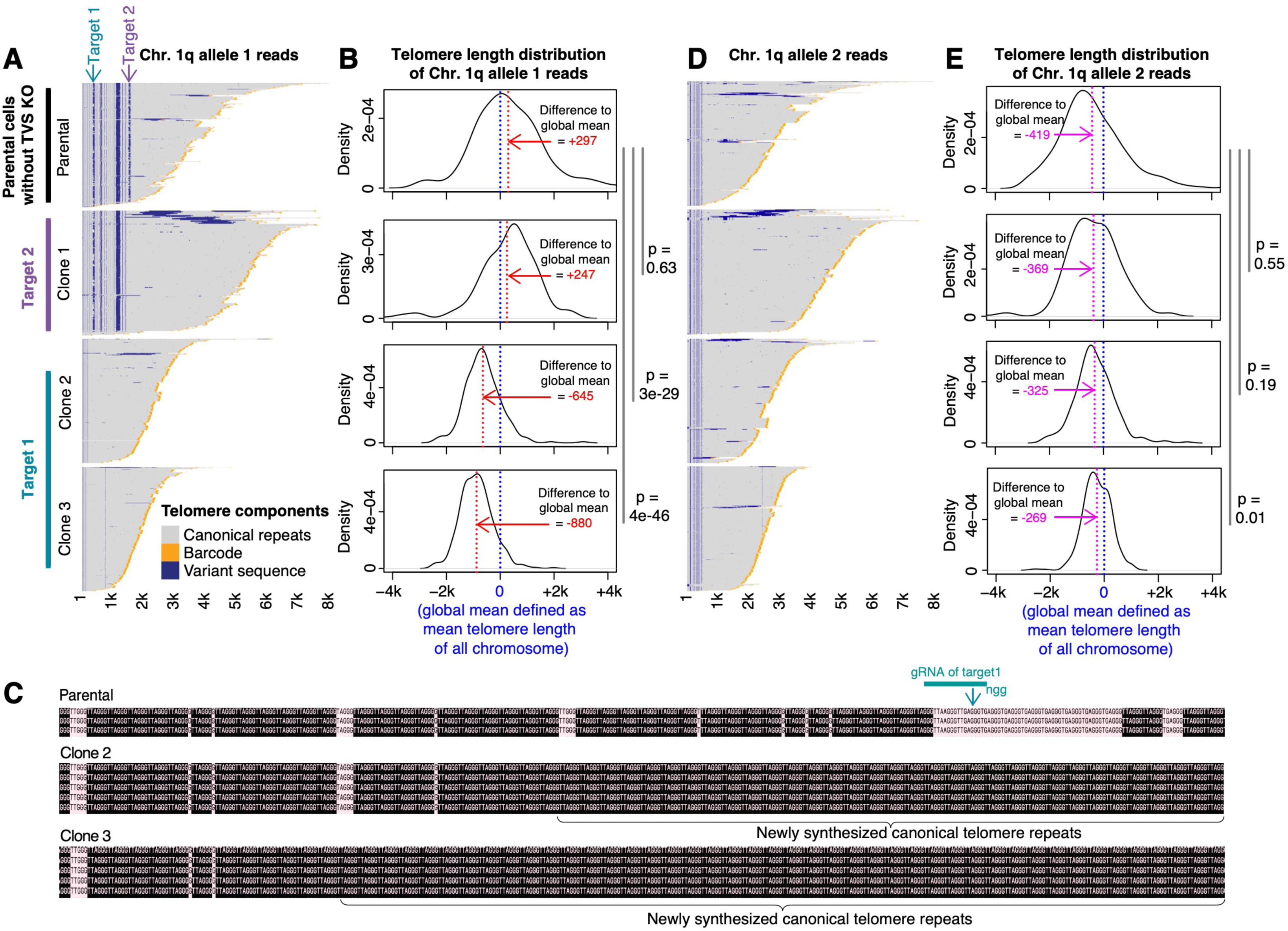
Allele-specific telomere length can be re-set by altering TVS abundance in clonal HCT116 cell lines. **A)** Telomere sequencing reads for the Chr. 1q allele 1 (Chr. 1q-A1) in parental HCT116 cells (top). Two CRISPR sgRNA target sites are indicated by the arrows on top. After TVS-specific knock out, three independent single-cell-derived clones were isolated, and the telomere sequencing reads from the edited Chr. 1q-A1 alleles are shown. Clone 1 harbors a deletion at the second sgRNA target, resulting in minimal changes in TVS abundance, whereas clones 2 and 3 harbor deletions at the first sgRNA target, resulting in the loss of most TVSs. **B)** Density plots of normalized telomere length for the Chr. 1q-A1 allele. To account for differences in mean telomere length between parental and edited clones, the mean telomere length across all chromosomes in each sample (blue dotted line) was subtracted from the raw Chr. 1q-A1 telomere length to yield a normalized telomere length for comparison. The changes of mean telomere length in Chr. 1q-A1 allele after TVSs knockout are shown (red dotted line). **C)** CRISPR sgRNA target 1 in the parental allele is shown as a segment (teal). Canonical telomere repeats (black), and variant sequences (pink) are shown around target 1. Newly synthesized canonical repeats replacing the deleted TVS blocks in clones 2 (middle) and 3 (bottom) are labelled. **D)** The telomere sequencing reads from the other non-targeted allele of Chr. 1q (Chr. 1q-A2). **E)** Density plots of normalized telomere length for Chr. 1q-A2. The telomere length relative to the global mean is shown (pink dotted line).

Given that each single-cell-derived clone had a different mean telomere length, we normalized the Chr. 1q-A1 telomere length in each clone to the global mean telomere length of that clone. The global mean telomere length in a clone is calculated as mean telomere length across all chromosomes within the clone. This normalization allowed direct comparison of allele-specific changes relative to overall telomere length (**Figure 5B**). As expected, deletion of the 3’ TVS-rich region at Target 1 led to a significant reduction in Chr. 1q-A1 telomere length by an average of 942-1177 bp in clones 2 and 3, compared to the parental line (p = 3e–29, and p = 4e–46, respectively; **Figure 5B**). By contrast, deletion at Target 2 had a negligible effect (p = 0.63, **Figure 5B**), confirming the importance of TVS abundance rather than simply CRISPR-induced editing.

As an internal control, we measured telomere length at the other non-targeted Chr. 1q allele 2 (Chr. 1q-A2) across all single-cell-derived clones. This allele remained unaffected in all cases (**Figures 5D–5E**), underscoring the allele-specific effect of the TVS deletion. Together, these results provide the first direct experimental evidence that TVS abundance and distribution actively regulate allele-specific telomere length.

## Discussion

The extreme heterogeneity in allele-specific telomere length revealed by recent long-read-sequencing data raise an important question on the underlying mechanism of allele-specific telomere maintenance. Our results indicate that the unique TVS distribution at each chromosomal end underlies allele-specific telomere length, which is both inheritable and relatively stable in offspring. The correlation becomes even more apparent in telomerase-positive cancer cells, where telomere length is actively maintained during continuous cell division. Over time, stochastic mutations introduced during DNA replication, particularly in germ cells and tissue stem/progenitor cells, contribute to the gradual evolution of allele-specific TVS patterns. Consequently, telomere length variation arises and is strongly associated with TVS abundance. In rare cases, recombination events between telomeres may occur during meiotic or mitotic divisions, leading to the exchange of genetic content between telomeres (**Supplementary Figures S6-S8**). Notably, such highly personalized and traceable features in telomere sequences are absent from current telomere-to-telomere (T2T) genome assemblies and are not captured in recently published human recombination maps (*45*).

Our attempt to associate allele-specific telomere length with the number of TRF1/TRF2 binding motifs, used as a proxy for telomere protein counting, yielded variable results (**Figure 3D**). Correlations were more robust in cells with shorter mean telomere lengths, such as HCT116 and the HT1080 B2 clone, but were weaker in telomerase-positive cells with longer telomeres, such as the HT1080 A6 and B3 clones. Several factors could explain this discrepancy. First, the telomere protein counting model is based on the binding of the entire shelterin complex, not just TRF1/TRF2, and the exact footprint and binding dynamics of shelterin components at the telomere remain poorly characterized. TVS interruptions may influence shelterin spacing, introducing uncertainty into binding site estimates based solely on TRF1/TRF2 motifs. Second, uncharacterized DNA-binding proteins in subtelomeric regions may modulate allele-specific shelterin recruitment, a possibility that warrants further investigation. Third, the telomere protein counting model was originally described in yeast, which has relatively short telomeres. An additional model based on replication fork progression, has recently been proposed (*16*). It is possible that both the telomere protein counting and replication fork progression models cooperate in telomere length regulation, especially in human cells with long telomeric tracks.

To probe these mechanisms further, emerging long-read sequencing techniques may be leveraged. For example, Fiber-seq (*46, 47*) has been used to investigate chromatin accessibility and CTCF binding in both telomeric and subtelomeric regions. Similarly, DiMeLo-seq (*48*) has proven effective in quantifying protein occupancy, such as CENP-A density, in highly repetitive centromeric regions. These approaches could be adapted to characterize shelterin binding density along individual telomeres, offering deeper insights into telomere maintenance at allele resolution.

Our data also reveal a clear asymmetry in telomere maintenance across generations, with lower stability observed during paternal transmission. While this difference may partly stem from the distinct DNA replication histories of oogenesis and spermatogenesis, differences in chromatin structure and epigenetic states could also play a role in shaping this parent-of-origin effect. For example, telomere maintenance dimorphism has been documented in animals of the order *Dasyuromorphia* (*49*), and a recent study of a four-generation human pedigree identified a strong paternal bias across all forms of germline de novo mutations (*50*). Moreover, parent-of-origin effects on telomere elongation associated with alternative lengthening of telomeres (ALT) have been observed as early as the two-cell stage in mouse embryos (*51*). Taken together, these findings support our conclusion that parent-of-origin effects may substantially influence the accumulation of telomeric variant sequences and contribute to telomere length heterogeneity.

It is well established that the length of the shortest telomeres, not the mean telomere length, predicts the viability and long-term proliferative capacity of tissue stem and progenitor cells (*52*). Given the allele-specific distribution of TVSs and the resulting variability in shelterin binding density, this feature may become especially important later in life. The larger and more abundant TVSs near subtelomeric regions may modulate telomere capping and play an important role in aging-related diseases. Our observation that each telomere allele harbors a unique TVS signature further underscores the individuality of telomere architecture. Accordingly, telomere length profiles are highly personal, and the specific alleles responsible for triggering senescence may differ among individuals during the aging process. Integrating clinical data from aging-related diseases with high-resolution characterization of uncapped telomeres in senescent cells may help identify telomere alleles that are particularly susceptible to dysfunction. In the future, the ability to detect and track such susceptible alleles may offer a more accurate predictor of aging-related disease risk than conventional measurements of average telomere length alone.

## Acknowledgements

This work was supported by a grant from the National Medical Research Council Singapore (MOH-CIRG23jan-0006) awarded to S.L.; The Japanese Society of Hematology Research Grant and JSPS Kakenhi Grants (23K24165 and 25K02519) to M. O.. J. Liu is supported by the National Natural Science Foundation of China (Grant No. 12371283), Shenzhen Fundamental Research Program (Grant No. JCYJ20240813113518024), 1+1+1 Collaborative Fund, and Shenzhen Stability Science Program. We thank Milton Kwek Sheng Yi and Jae Peck Yean Tan (Research Instruments Pte Ltd) for their assistance with Oxford Nanopore sequencing. We also thank Qi Yang Li, Eric Yang Shun Kai, and John Zhang (NovogeneAIT) for their support with PacBio HiFi sequencing and Nanopore ultra-long sequencing.

## Supplementary Data

**Figure S1:**
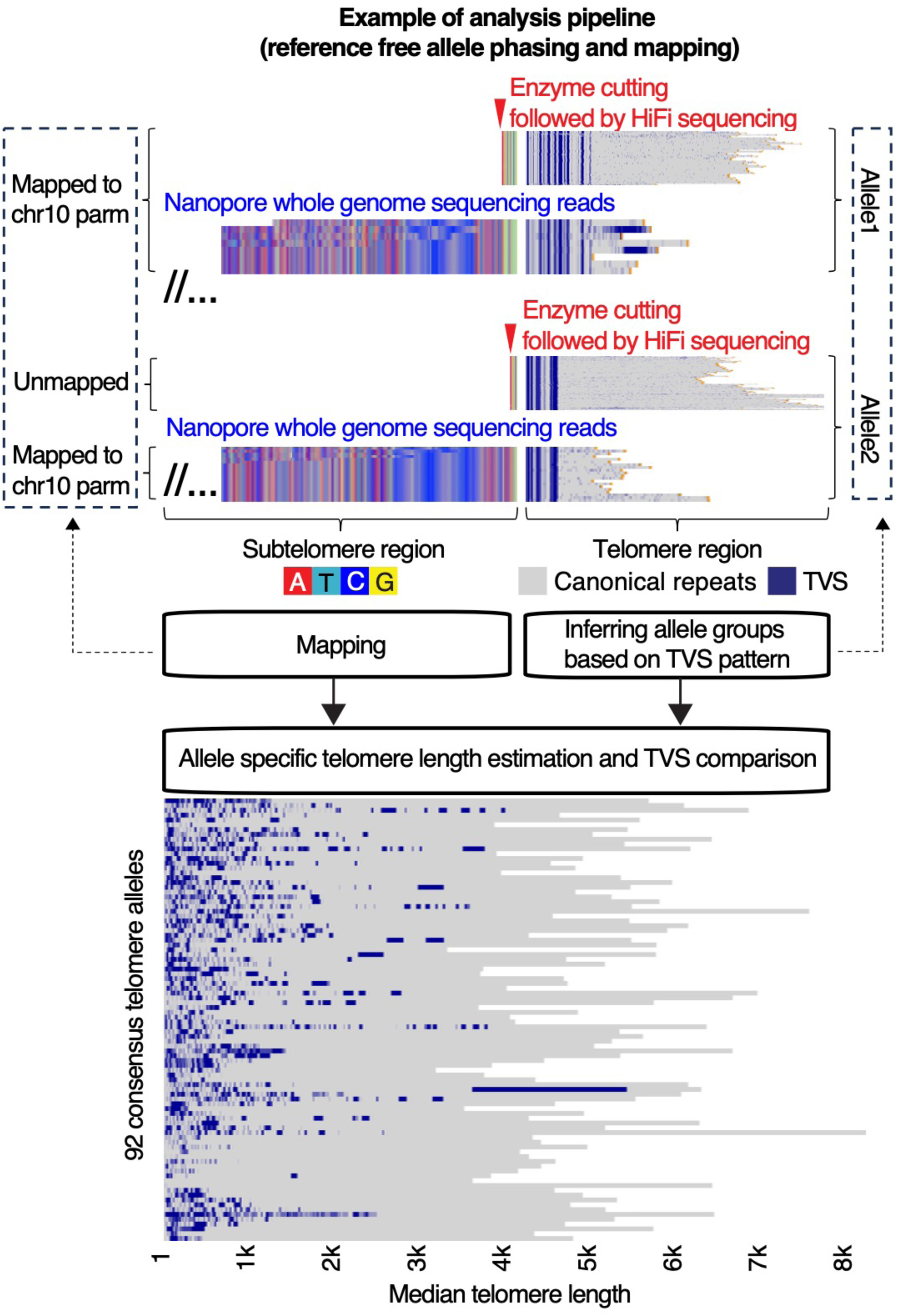
Analysis pipeline for reference-free allele phasing and chromosome assignment. Telomere start sites were defined as the first occurrence of the sequence 5′-TTAGGGTTAGGG-3′. Each sequencing read was split into two parts: the telomeric region and the upstream subtelomeric region. Two sets of long-read data—PacBio HiFi and Oxford Nanopore—were collected from both telomere-enriched and whole-genome sequencing (WGS) libraries. Telomere-containing reads from both sources were independently phased based on their TVS patterns. Within each allele, reads from the telomere-enriched and WGS datasets could be matched via their unique, allele-specific TVS profiles. In cases where mapping of telomere-enriched reads was not possible due to limited subtelomeric sequence, chromosomal assignment was inferred from the corresponding WGS reads (Nanopore or PacBio Revio). This approach enabled the identification and chromosomal assignment of nearly all 92 allele groups within each sample.

**Figure S2:**
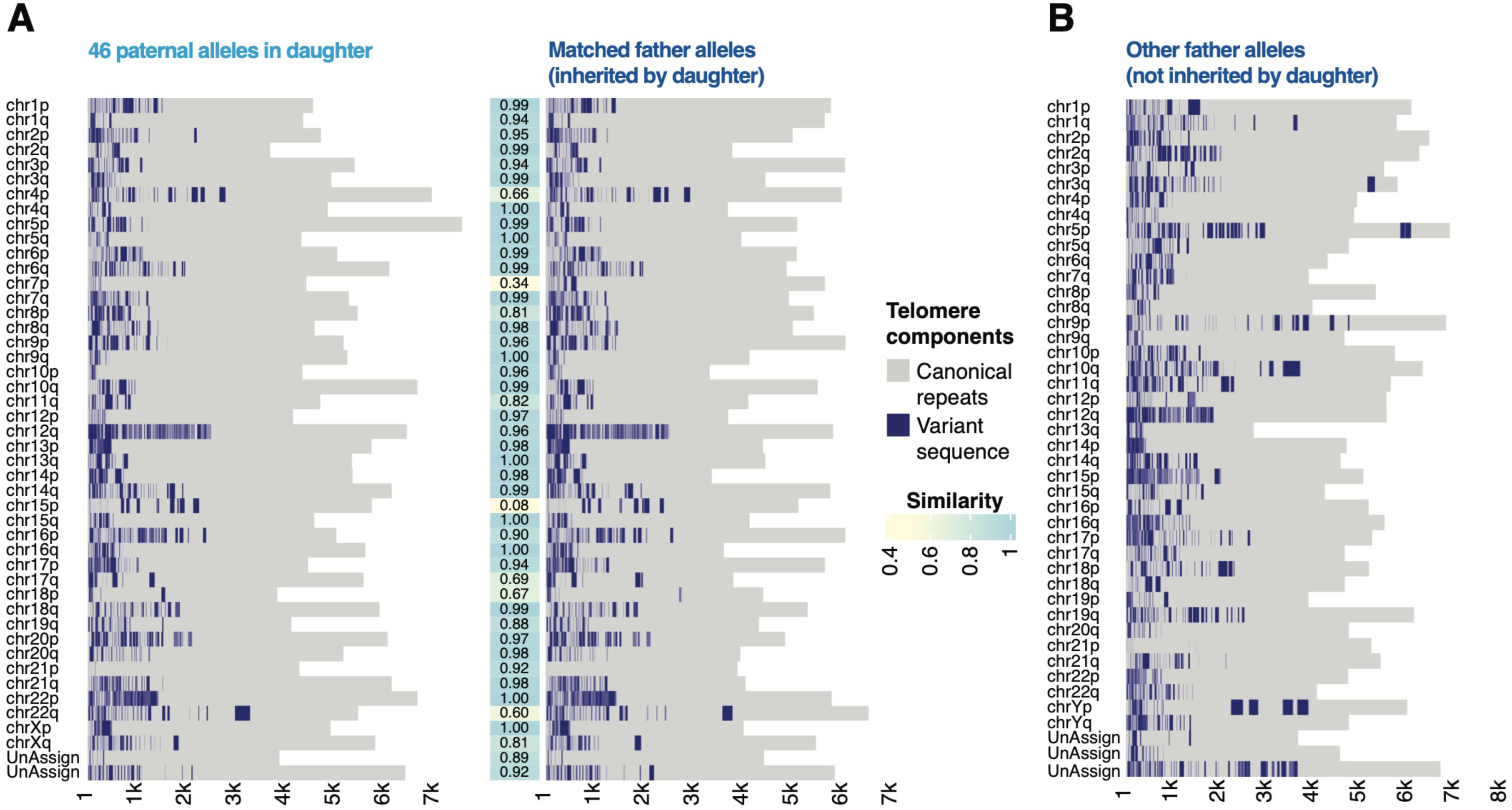
Paternal allele-specific telomeres and inheritance in family 1. **A)** The distribution of the 46 telomere clusters/alleles in the father’s peripheral blood leukocytes that are inherited by the daughter (see also Figure 1C). **B)** The remaining 44 paternal allele-specific telomeres not inherited by the daughter, completing the full set of 90 paternal alleles.

**Figure S3:**
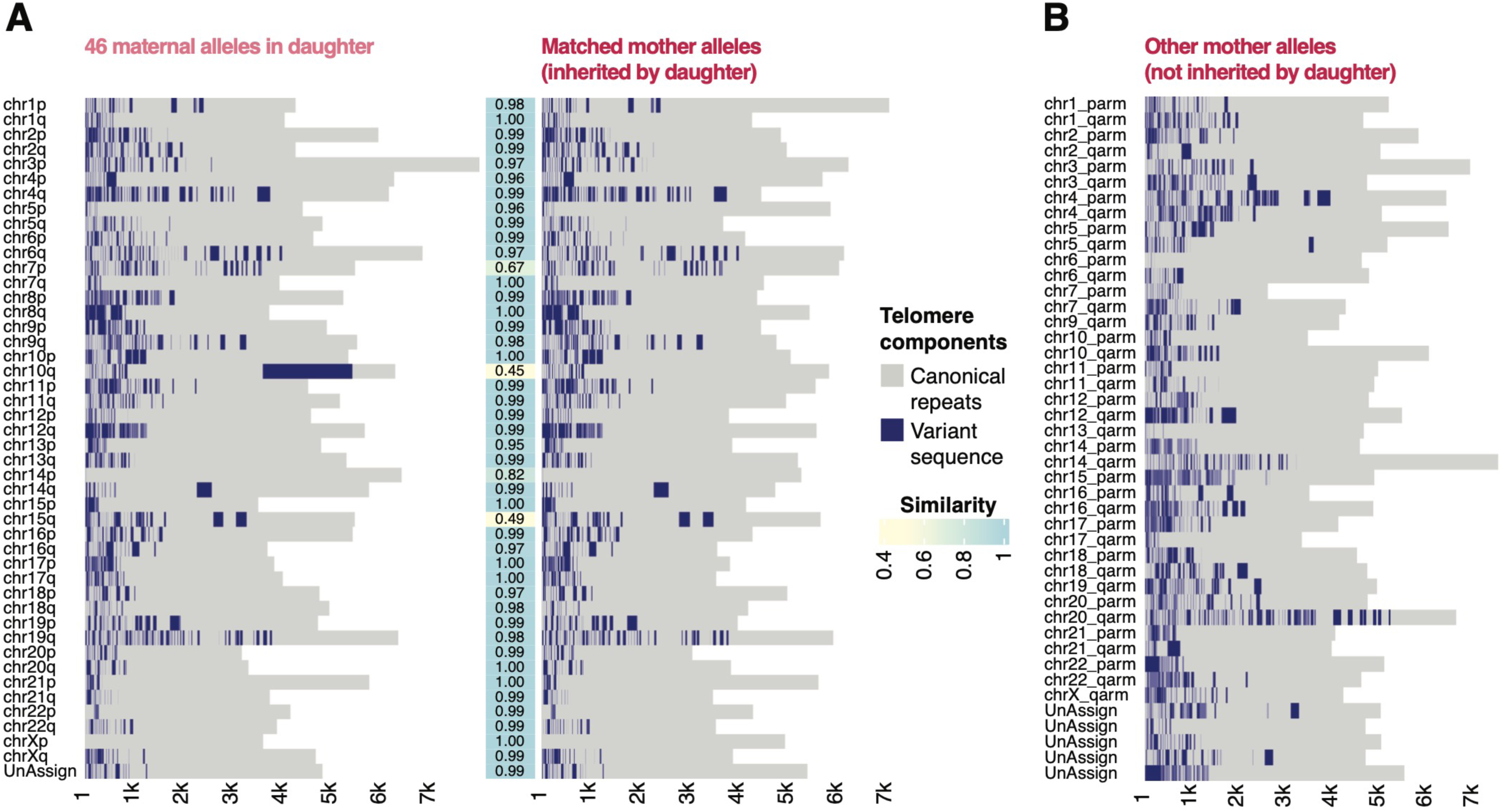
Maternal allele-specific telomeres and inheritance in family 1. **A)** The distribution of the 46 telomere clusters/alleles in the mother’s peripheral blood leukocytes that are inherited by the daughter (see also Figure 1C). **B)** The remaining 44 maternal allele-specific telomeres not inherited by the daughter, completing the full set of 90 maternal alleles.

**Figure S4:**
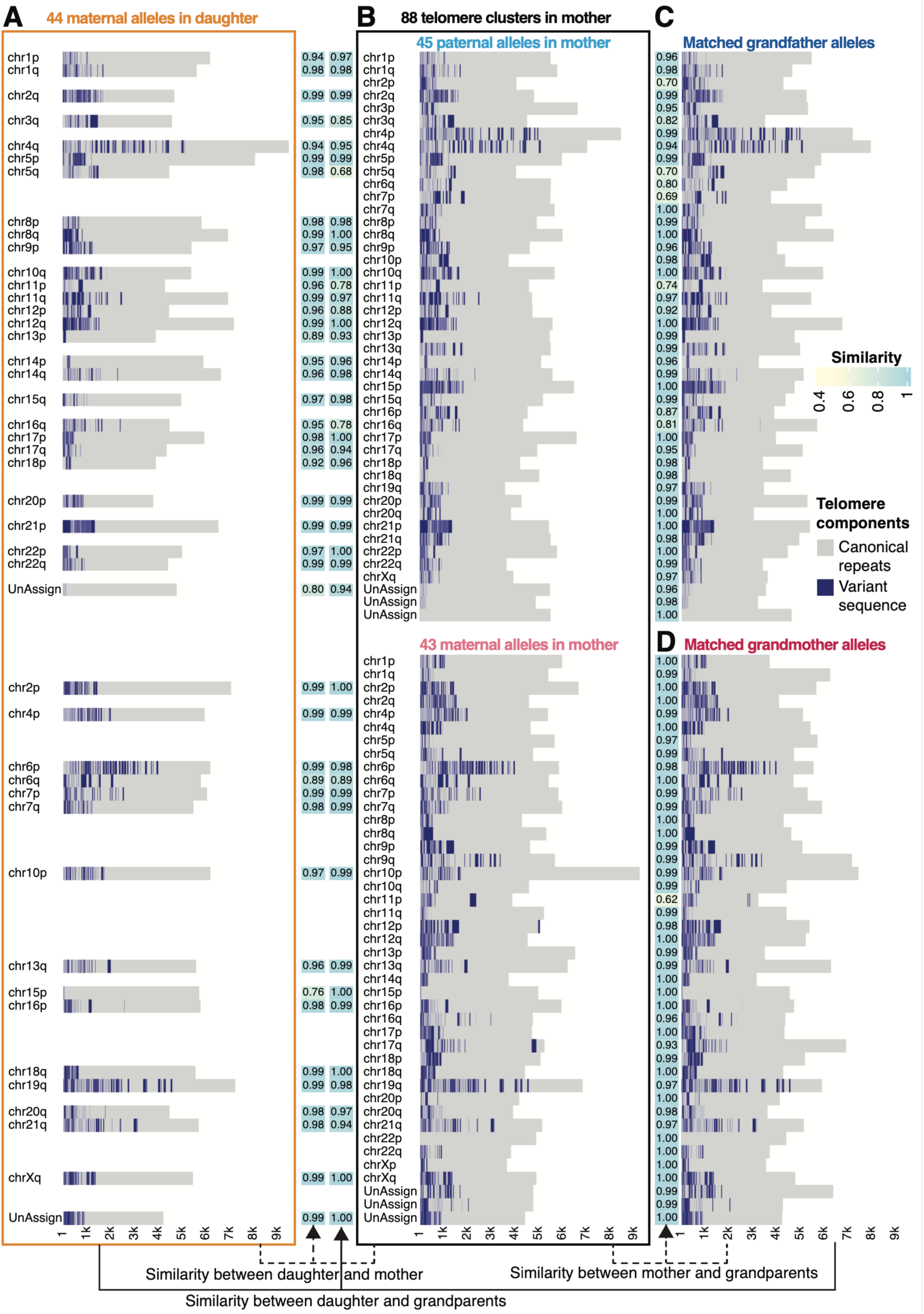
Stable and inheritable allele-specific telomere variant sequences (TVSs) in family 2. **A)** Distribution of TVSs in 44 telomere clusters/alleles in the daughter’s peripheral blood leukocytes (PBLs). **B-D)** Full set of 88 telomere clusters/alleles identified in the mother’s PBLs, with 45 alleles traced back to the grandfather C) and 43 to the grandmother D). Pairwise similarity scores between alleles are visualized as heatmaps between the respective individuals.

**Figure S5:**
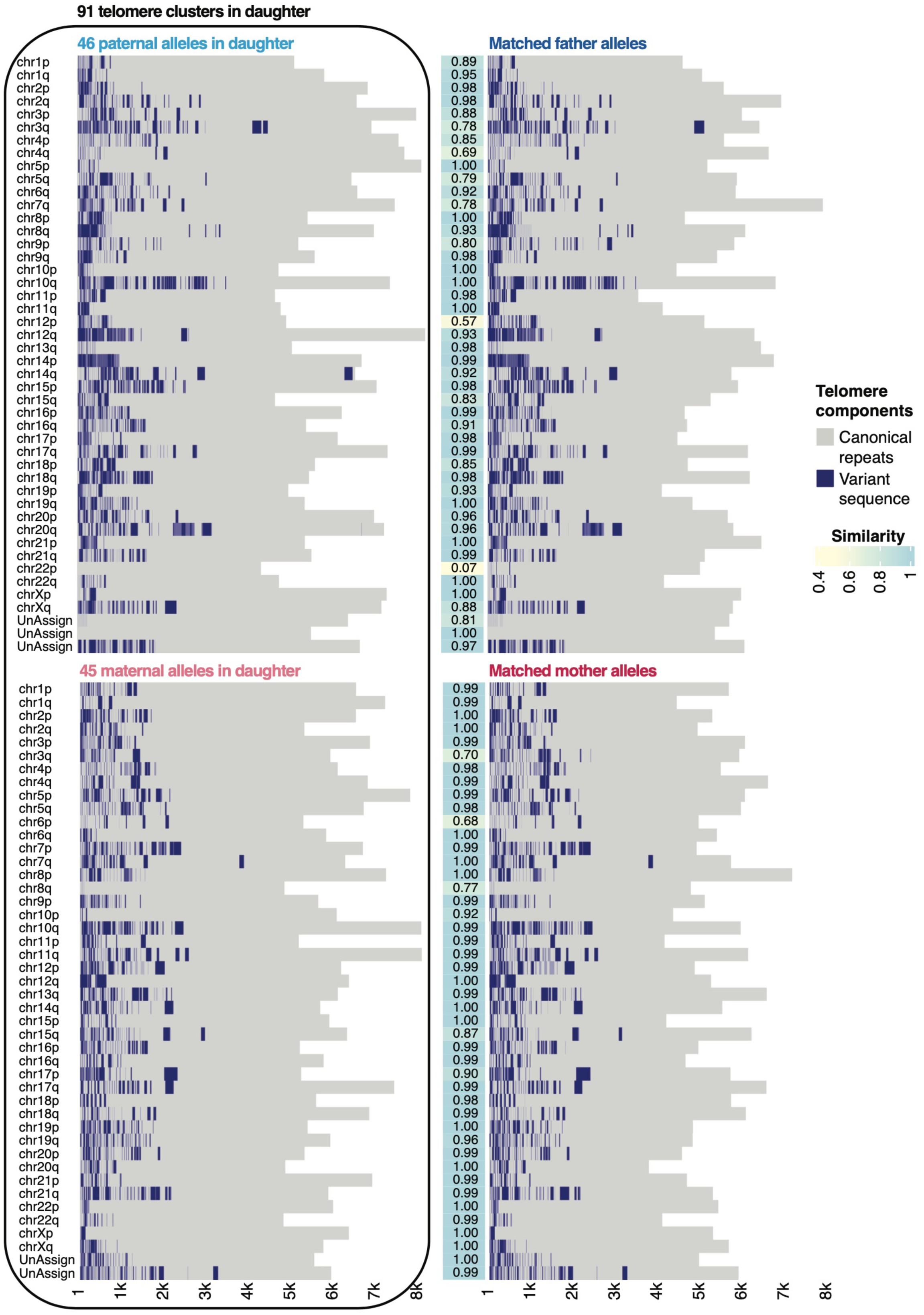
Stable and inheritable allele-specific telomere variant sequences (TVSs) in family 3. Distribution of TVSs across 91 allele-specific telomeres identified in the daughter’s peripheral blood leukocytes, including 46 paternal and 45 maternal alleles (left), and the corresponding TVS-matched alleles from the father and mother (right) panel. Allele similarity is presented as a heatmap (middle).

**Figure S6:**
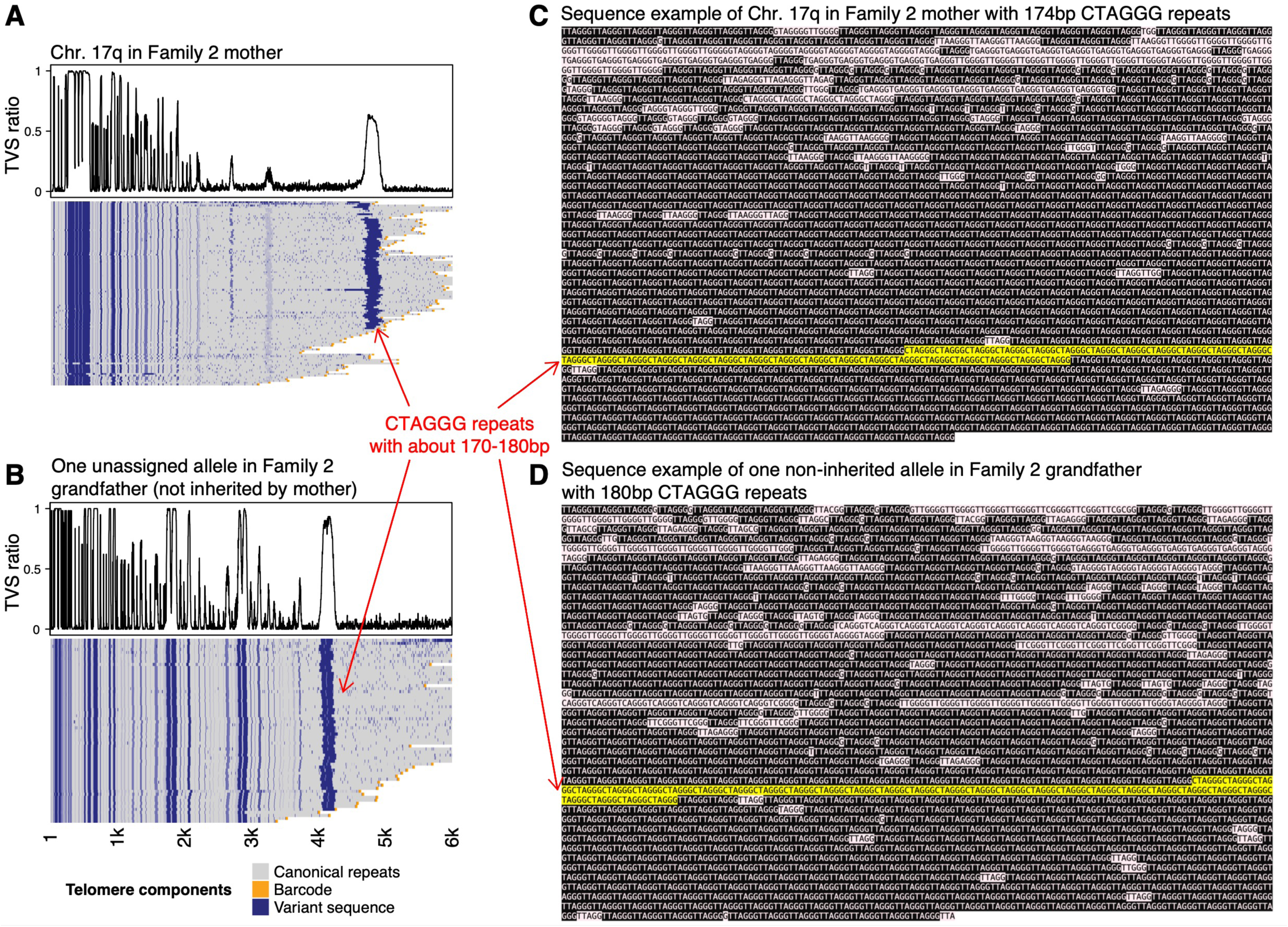
Recombination event on Chr. 17q identified in the mother from family 2. **A)** Distribution of telomere variant sequences (TVSs) along Chr. 17q in the mother. The terminal TVS block consists of ∼170–180 bp of CTAGGG repeats. **B)** An almost identical CTAGGG repeat block (∼170–180 bp) was observed in only one of the grandfather’s telomere alleles, which was not inherited by the mother. **C**–**D)** Representative sequences from the mother’s Chr. 17q and the potential grandfather allele involved in the recombination event. Canonical telomeric repeats (black), variant sequences (pink), and the putative recombined TVS block (yellow) are shown.

**Figure S7:**
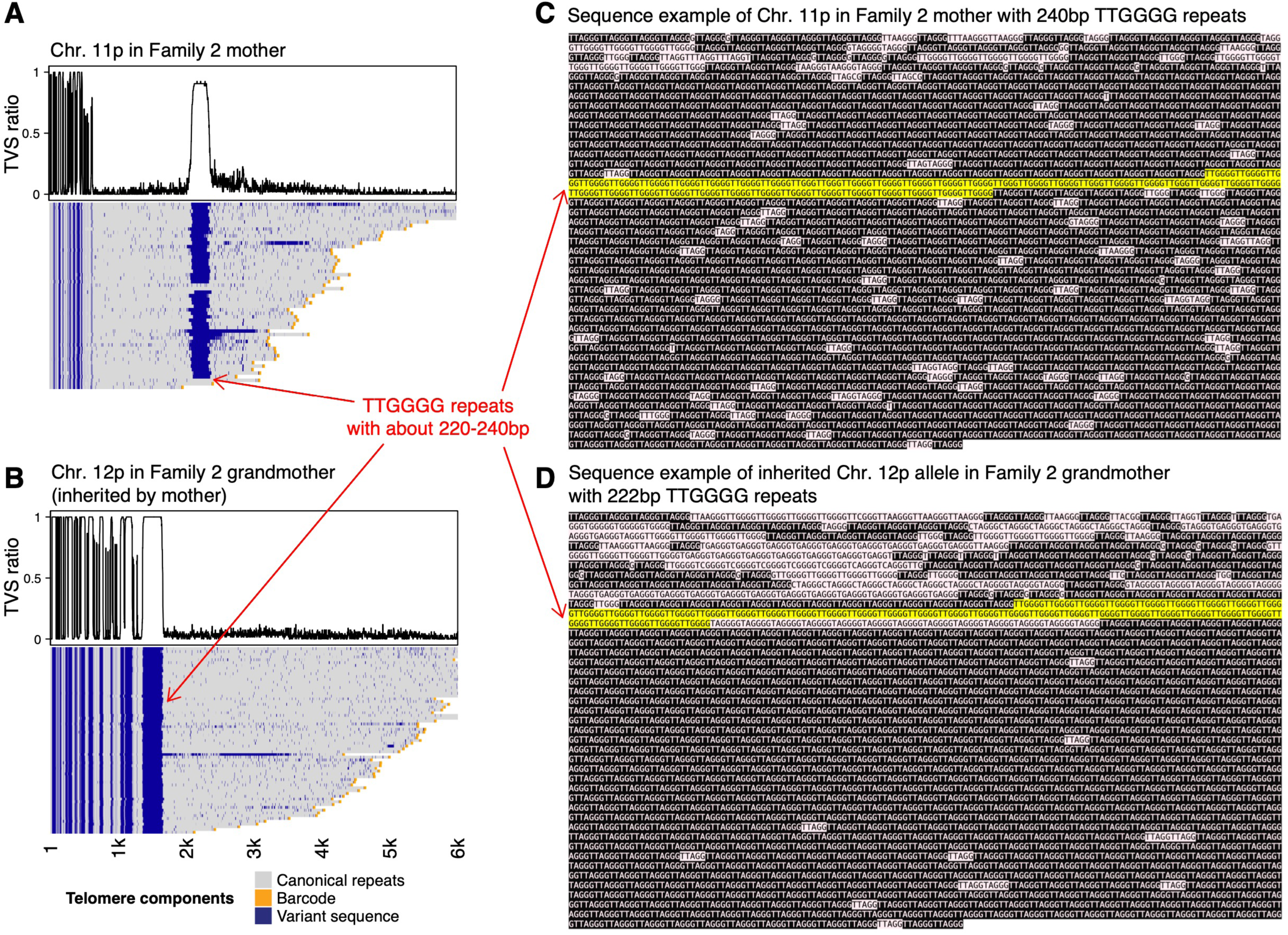
Chromosome exchange event on Chr. 11p identified in the mother from family 2. **A)** Distribution of TVSs along Chr. 11p in the mother. The terminal TVS block consists of ∼220–240 bp of TTGGGG repeats. **B)** An almost identical TTGGGG repeat block (∼200–240 bp) was observed in one of the grandmother’s telomere alleles, which was inherited by the mother. **C**–**D)** Representative sequences from the mother’s Chr. 11p and the putative donor sequence on Chr. 12p involved in the gene conversion event. Canonical telomeric repeats are (black), variant sequences (pink), and the putative gene-converted TVS block (yellow) are shown.

**Figure S8:**
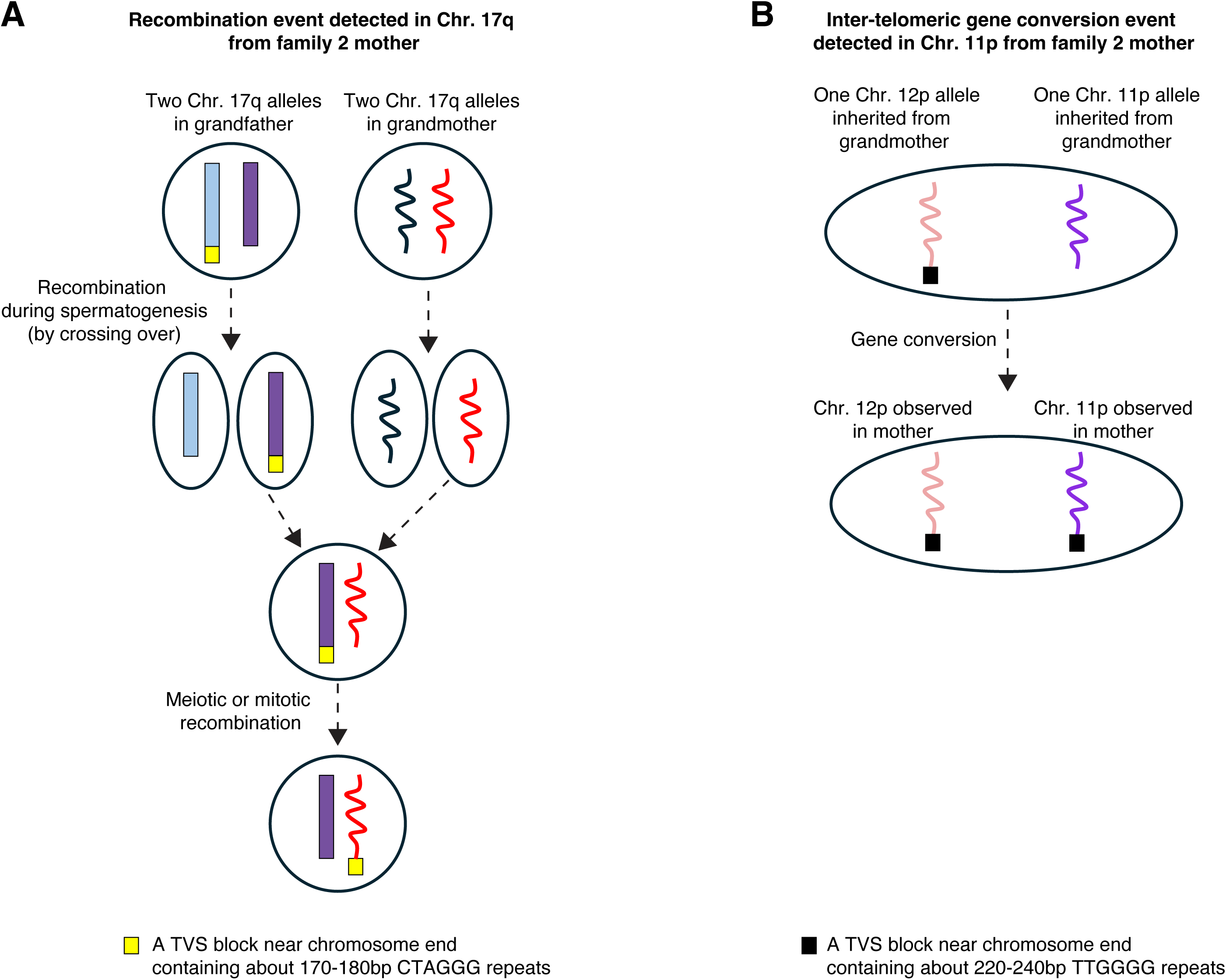
Schematic of recombination and gene conversion events identified in family 2. **A)** Schematic of the Chr. 17q recombination event observed in the mother, likely involving a non-inherited allele from the grandfather. **B)** Schematic of the Chr. 11p gene conversion event in the mother, likely involving an inherited allele from the grandmother and potentially involving Chr. 12p.

**Figure S9:**
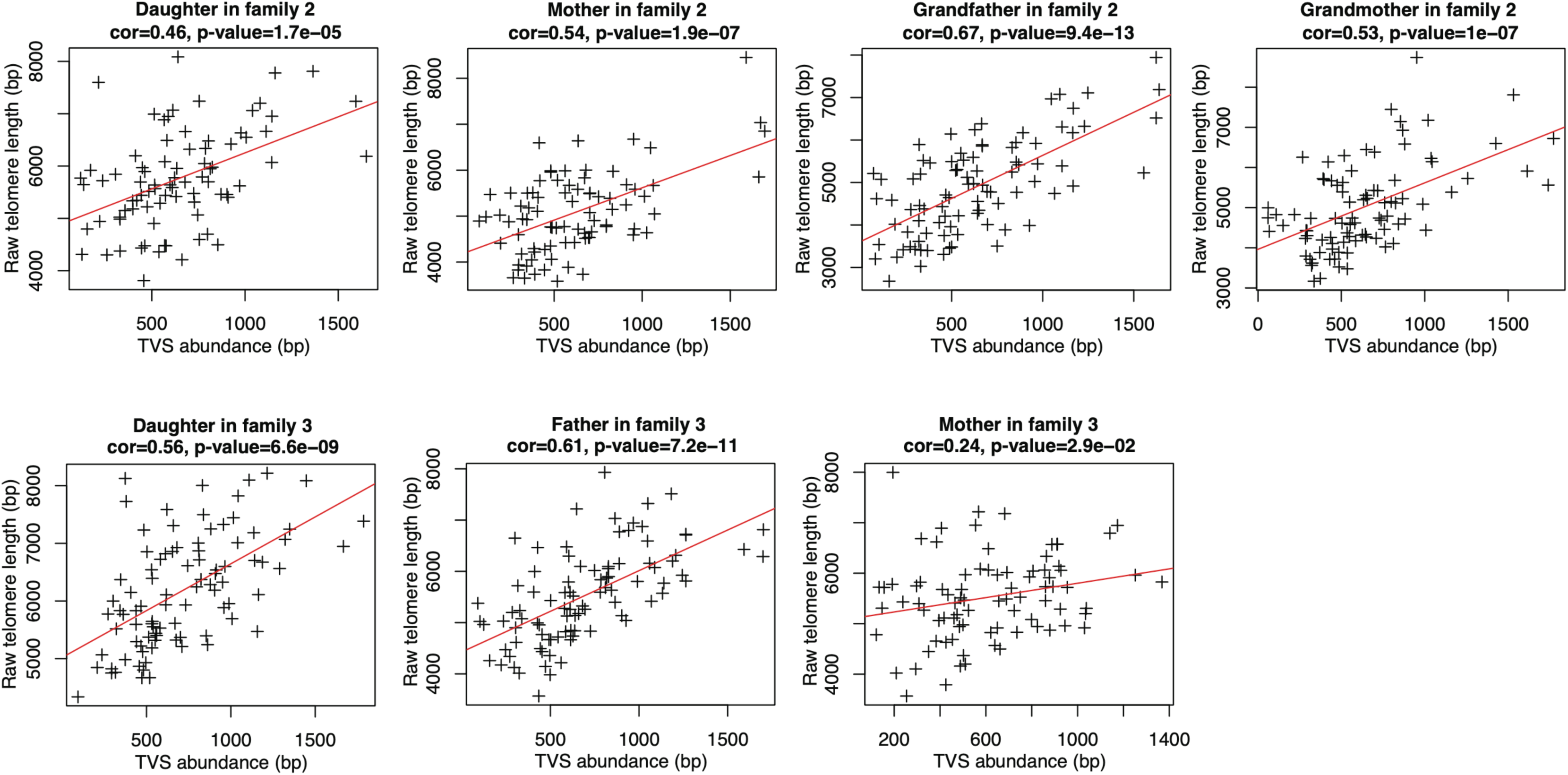
Positive correlation between TVS abundance and telomere length in PBLs of families 2 and 3. Scatter plots showing the relationship between allele-specific telomere length and TVS abundance for each individual in families 2 and 3. The linear regression fit for each dataset is shown (red line).

**Figure S10:**
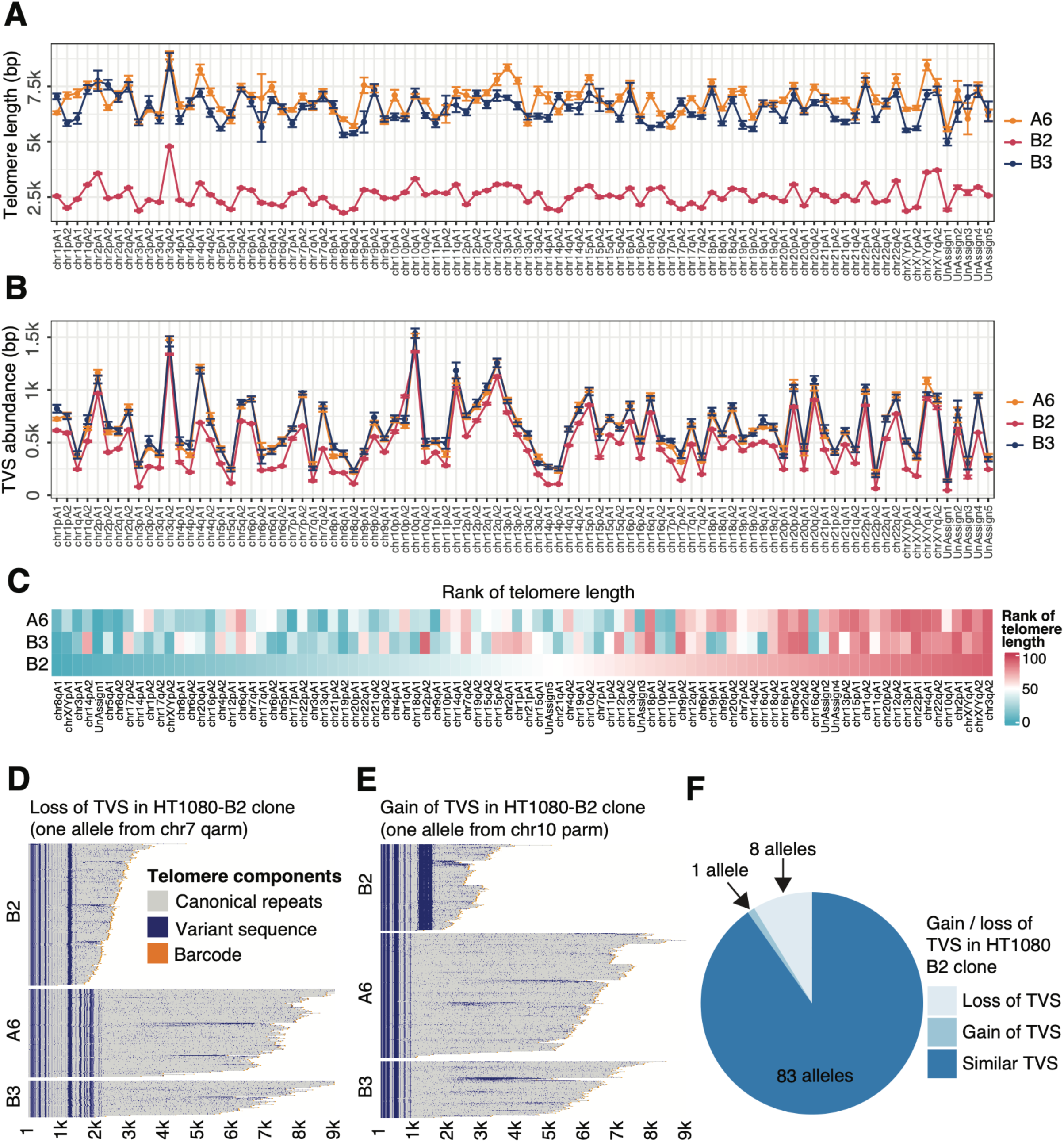
Comparison of allele-specific telomere length and TVS distribution in three single-cell-derived HT1080 subclones. **A**–**B)** Allele-specific telomere length distributions and TVS abundance profiles across three single-cell-derived HT1080 clones (HT1080-A6, HT1080-B3, and HT1080-B2). **C)** Heatmap showing the rank order of allele-specific telomere lengths across the three subclones **D)** Example of TVS loss in the HT1080-B2 clone, indicative of telomere re-setting. **E)** Example of TVS gain in HT1080-B2. **F)** Percentage of telomere alleles showing gain or loss of TSVs compared to A6 and B3.

**Figure S11:**
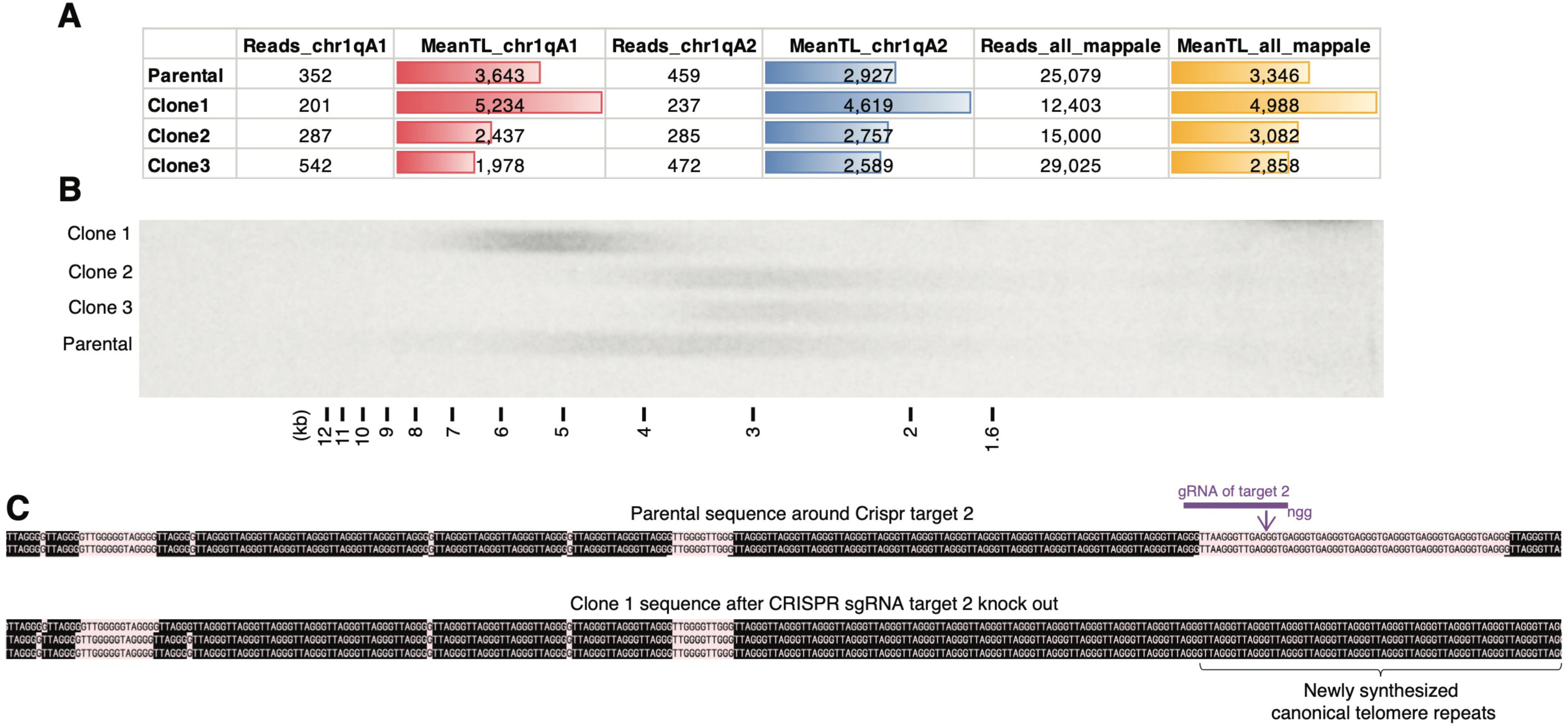
Experimental validation of TVS knockout and telomere length effects in single-cell-derived HCT116 subclones. **A)** Summary of read counts and mean telomere lengths for the two Chr. 1q alleles and all mapped telomere reads in HCT116 parental cells and subclones 1–3. Clones 2 and 3 harbor large TVS deletions introduced via CRISPR targeting site 1, while clone 1 carries a smaller deletion via CRISPR targeting site 2 (see also Figure 5). **B)** Teloblot analysis showing telomere length distributions in HCT116 parental cells and three subclones following TVS knockout. **C)** Sequence comparison around CRISPR target 2 in parental HCT116 cells and clone 1. Canonical telomeric repeats (black) and variant sequences (pink) are shown.

**Extended Data 1: Detailed telomere reads in Family 1.** Panels 1–92 display the individual telomere reads for each of the daughter’s 92 telomere alleles, alongside the corresponding parental alleles with matched TVS patterns.

**Extended Data 2: Detailed telomere reads in family 2.** Panels 1–44 display the individual telomere reads for each of the daughter’s 44 assigned telomere alleles, alongside the corresponding TVS-matched alleles from her mother, grandfather (GF), and grandmother (GM).

## Methods

### Cell lines and Plasmid

Parental HCT116 and HT1080 cell lines (ATCC) were maintained in DMEM supplemented with 10% fetal bovine serum (FBS) and 1% Penicillin-Streptomycin and cultured in a humidified incubator with 5% CO_2_ at 37°C. Cells were sub-cultured at a 1:8 ratio and seeded at a density of 5 × 10⁵ cells per 10 cm dish for experiments.

### Human subjects and sample processing

The subjects from Japan were recruited at the Misakaenosono Mutsumi Developmental, Medical and Welfare Center, for a research project entitled “Telomere shortening in chr. 21 trisomy patients”. Written informed consent was obtained from participants upon enrolment. Kumamoto University Institutional Review Board had approved the study protocol (genomic research No. 557). The subjects from Singapore were recruited from the Cardiac Ageing Study (CAS)(*53*), a prospective study that examines characteristics and determinants of cardiovascular function in older adults. Written informed consent was obtained from participants upon enrolment. The SingHealth Centralised Institutional Review Board had approved the study protocol (CIRC/2014/628/C).

Antecubital venous blood samples were collected into PAXgene Blood DNA tubes (BD Biosciences). For fresh samples, processing was carried out immediately after collection. For archived samples, blood was placed on ice for transportat and processed within 6 hours to isolate buffy coat, which was then stored at −80 °C. Genomic DNA from PBLs was extracted using the PAXgene Blood DNA Kit (Qiagen) according to the manufactuerer’s conditions.

### Genomic DNA extraction and Southern blotting (Teloblot)

Genomic DNA from cultured cells was extracted using the Gentra Puregene Genomic DNA Purification Kit (Qiagen). Telomere Southern hybridization was performed as previously described (*54*). Briefly, 0.5 µg of genomic DNA was digested with RsaI/HinfI or EcoRI (as indicated) at 37°C for 16 hours. The digested DNA was normalized and resolved on a 0.6% agarose gel (SeaKem ME Agarose, Lonza) in 1x TBE buffer. Following electrophoresis, the gel was processed through a series of treatments with gentle shaking at room temperature: depurination in 0.25 M HCl for 30 minutes, denaturation in 0.5M NaOH and 1.5M NaCl for 30 minutes, and neutralization buffer (1M Tris pH 7.4, 1.5M NaCl) for 30 minutes. DNA was then transferred overnight by capillary blotting onto a Hybond-XL membrane (GE Healthcare) using 10x SSC buffer and Whatman GB003 gel blot paper (GE Healthcare). After transfer, the membrane was UV-crosslinked using an XL-1500 UV crosslinker (Krackeler Scientific Inc.) and prehybridized in phosphate buffer (0.5M NaPO4, pH 7.2, 7% SDS, 1mM EDTA, pH 8.0) for 1 hour at 37 °C with rotation. The membrane was then hybridized overnight with a ³²P-labeled (TTAGGG)₆ oligonucleotide probe in the same buffer at 37 °C with rotation. Post-hybridization washes were performed three times for 15 minutes each in phosphate wash buffer (0.2M NaPO4 pH 7.2, 1%SDS,1mM EDTA) at 25 °C. The membrane was then exposed to a phosphor screen for imaging.

### Data pre-processing of telomere-enriched sequencing reads and WGS reads

Telomere-enriched sequencing was performed on the PacBio Sequel IIe platform using polymerase 2.2. Consensus HiFi reads were generated on-instrument using the default CCS function. Genomic DNA without telomere enrichment was subjected to whole genome sequencing (WGS) using either the PacBio Revio or Oxford Nanopore platform.

For PacBio HiFi sequencing, telomere-containing reads were identified as those containing a sample barcode at the 3’ end and at least two consecutive canonical telomere repeats (“TTAGGGTTAGGG”). Sample demultiplexing was performed using lima (v2.6.99, part of the SMRT Link v11 package). Telomere repeat motifs were detected using seqkit (v2.4.0) (*55*) without allowing mismatches. The subtelomere–telomere boundary was defined as the first occurrence of two consecutive canonical telomere repeats in the 5’ to 3’ direction. After demultiplexing, mean telomere length for each sample was computed as the mean distance between the telomere start site and the barcode.

Nanopore sequencing was carried out on the PromethION 2 Solo platform using an R10.4.1 flow cell (FLO-PRO114M) and sequencing kit SQK-LSK114. Iinitial base calling was performed on-device with MinKNOW v25.05.14 (https://nanoporetech.com/software/devices/p2-solo/software) using the “fast” model. To maximize read accuracy, a second base calling step was applied to reads labeled “pass” in the initial output using software dorado-0.9.5 (https://github.com/nanoporetech/dorado) with the “super accurate” mode. Further read correction was performed using the HERRO pipeline (https://github.com/lbcb-sci/herro). WGS reads generated from the PacBio Revio platform were processed using the same strategy as Nanopore reads, except requiring two rounds of base calling and read correction. The resulting WGS reads from both platforms were pooled for downstream telomere analysis.

Telomere-containing reads in the WGS datasets were selected using the same criteria as for HiFi reads, except that no barcode was required. To distinguish genuine telomeric sequences from randomly distributed interstitial telomeric sequences, a two-step definition for telomere start site detection was implemented for WGS reads. First, a contiguous telomeric region was defined as a block in which the distance between any two canonical telomeric repeat units did not exceed 1,500bp. Within such validated block, the telomere start site was defined as the first occurrence of two consecutive canonical telomere repeats, in the 5’ to 3’ direction.

### Construction of customized reference genome and mapping

A customized reference genome that incorporates both the entire T2T genome (CHM13v2.0) and all chromosome-end contigs was constructed from 94 recently published pangenome assemblies. The CHM13v2.0 genome was obtained from the T2T Consortium website (https://github.com/marbl/CHM13), while pangenome sequences were downloaded from the Human Pangenome Project Freeze1-v2 release (https://github.com/human-pangenomics/HPP_Year1_Assemblies), including both all contigs and the subset of contigs mappable to the CHM13 genome. The full CHM13 genome was retained and only those pangenome contigs that mapped to either the first or last 250kb of any CHM13 chromosome were appended, thereby enriching the reference for subtelomeric diversity. Only the subtelomeric regions of WGS reads were used for alignment to minimize artifacts arising from telomere length variability and interspersed TVSs. Mapping was performed using Winnowmap v2.03 (*56, 57*), with the*-ax map-pb* setting for PacBio HiFi reads and *-ax map-ont* for Nanopore reads.

### Allele group identification based on telomere enriched reads from PacBio HiFi sequencing

Both the TVS pattern and the subtelomeric sequences near the subtelomere/telomere boundary were used to assess the similarity between telomere-enriched reads. Specifically, each telomeric region was converted into a numerical vector: canonical repeats were encoded as ‘1’, barcode sequences as ‘2’, and TVSs interspersed among canonical repeats as ‘3’. Similarly, the subtelomeric region was encoded as a nucleotide vector, where ‘0’, ‘1’, ‘2’, and ‘3’ represented A, T, C, and G, respectively.

To construct a unified representation of each read, the last 200bp of the subtelomeric vector and the first 2000bp of the telomeric vector were concatenated to form a single numeric vector. Pairwise similarity between reads was then computed as a correlation matrix derived from these concatenated vectors.

Dimensionality reduction was performed using t-distributed stochastic neighbor embedding (t-SNE) via the Rtsne R package (v0.17) (*41*), using the top two components for downstream clustering. To identify allele groups while accounting for noise, we applied HDBSCAN using the dbscan R package (v1.2.2). The minimum cluster size threshold was manually adjusted to optimize separation and ensure accurate detection of all 92 allele groups per sample.

### Construction of consensus alleles and allele-specific TVS ratio/signature

To represent the general TVS pattern within an allele group, a consensus allele was created by analyzing the TVS distribution at each base position along the telomeric region. Each base was defined as a TVS position if more than 50% of the reads in the allele group contained a TVS at that location; otherwise, the base was considered part of a canonical repeat. The length of the consensus allele was set to the median read length of all reads within the allele group.

In parallel, the allele-specific TVS ratio was designed to quantify the TVS distribution along the telomere without applying a binary classification. For each position, the TVS ratio was calculated as the percentage of reads containing a TVS at that position. The resulting TVS signature—a vector of positional TVS ratios starting from the telomere start site—captures the variability in TVS frequency across the telomere and can be visualized as a line plot.

### Chromosome assignment based on WGS sequencing

The same allele clustering strategy used for telomere-enriched reads was concurrently applied to telomere-containing reads from WGS, generating a corresponding set of phased allele groups and their positional TVS ratio profiles. To establish a one-to-one correspondence between the WGS-derived clusters and the telomere-enriched clusters, a correlation analysis was performed. Specifically, the TVS ratio profile of each WGS allele group was compared with all telomere-enriched allele groups by calculating pairwise correlation scores. Each WGS cluster was then linked to the telomere-enriched cluster with which it showed the highest positive correlation. Once this correspondence was established, the chromosome assignment from the WGS clusters could be directly transferred to the matched telomere-enriched clusters derived from PacBio HiFi reads.

### Merging of different sets of telomere-enriched reads

For all family samples and the HT1080 cell line, telomere enrichment was performed using an enzyme combination of RsaI, HinfI, and EcoRI. For the HCT116 cell line, two distinct enrichment strategies were applied in parallel to ensure comprehensive TVS capture across all telomere alleles. The first strategy used the RsaI-based combination (RsaI/HinfI/EcoRI), while the second employed SmiI-based combinations (BseJI/SmiI, Eco32I/SmiI and StuI/SmiI). To create a unified dataset for HCT116, allele groups inferred from the two enrichment methods were merged using a correlation-based strategy similar to that used for chromosome assignment. Specifically, pairwise correlation scores were calculated between the TVS ratio profiles of allele groups derived from each method, and the pair with the highest positive correlation was considered a matched group for integration.

Additional barcodes used for SmiI based telomere enrichment are shown below:

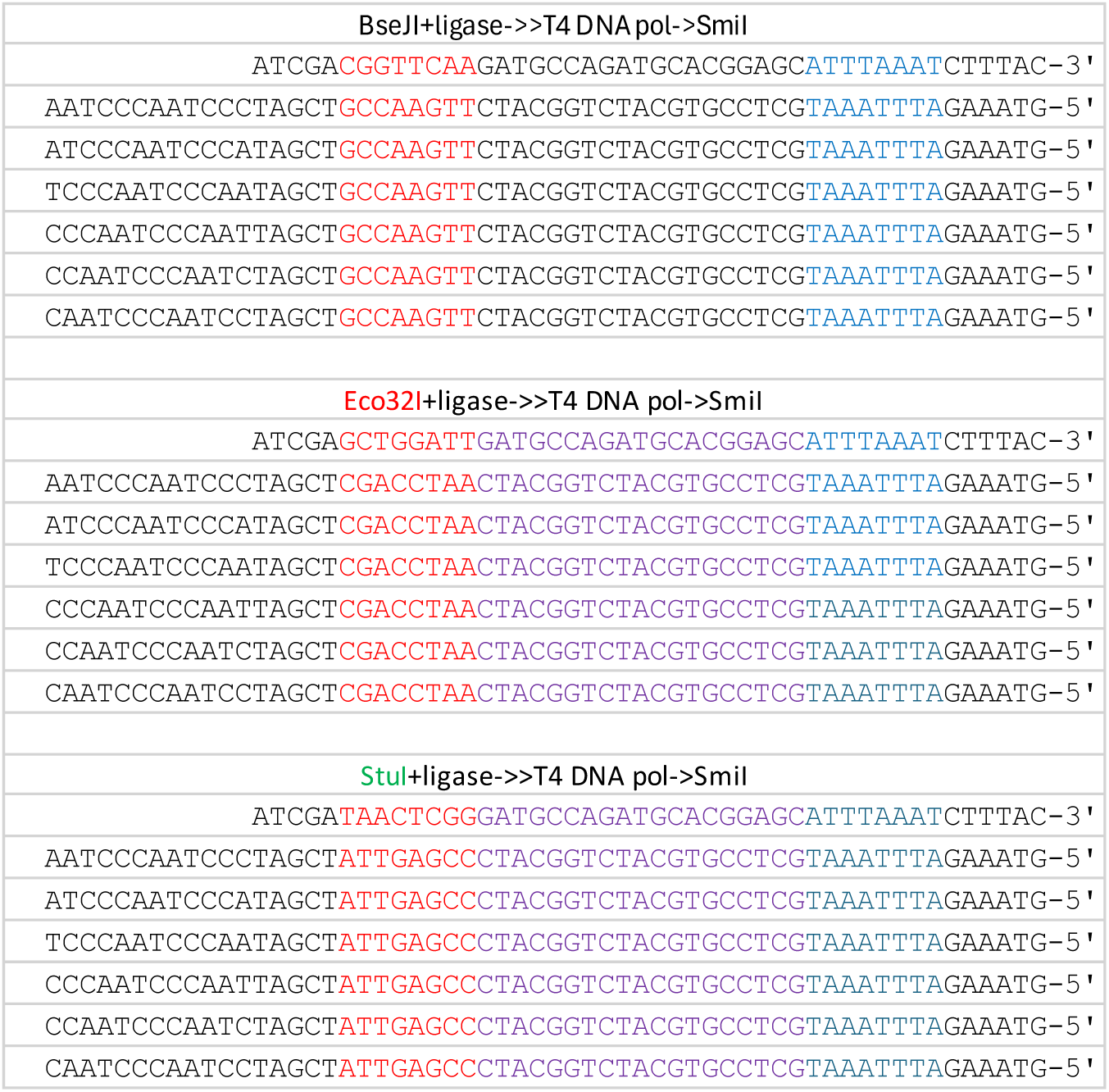

### Association between telomere length and TVS abundance

The association between raw telomere length and TVS abundance was calculated from consensus alleles. For each sample, Pearson correlation coefficients and corresponding p-values were calculated to quantify the relationship between TVS abundance (i.e., the number of TVSs observed in each consensus allele) and telomere length (defined as the median read length within each allele group). Only allele groups supported by more than 20 reads were included in this analysis. Linear regression was performed to visualize the trend and is plotted as a fitted line.

### Calculation of TRF1/TRF2 binding sites in telomere-enriched reads

Due to the higher sequencing accuracy of PacBio HiFi reads (Q33, as reported by PacBio’s: https://www.pacb.com/technology/hifi-sequencing/) compared to Nanopore WGS reads (Q21–26, as reported by Oxford Nanopore: https://nanoporetech.com/platform/accuracy), the analysis of TRF1/TRF2 binding motifs was restricted to the telomere-enriched PacBio HiFi reads. Binding sites were identified by searching for non-overlapping instances of the motif sequence “TTAGGGTTA” within the telomeric region of each read.

### Family stratification based on TVS similarity

To assess family-level relationships based on TVS patterns, we first constructed a distance matrix using TVS similarity. In this analysis, the three daughters (one from each family) were used as anchor samples. All allele clusters identified in these daughters were pooled and treated as unique reference features. Specifically, the daughter from Family 1 had 92 clusters, Family 2 had 90, and Family 3 had 91, resulting in 273 unique features in total.

For each individual sample, we compared every unique feature (i.e., each daughter’s allele cluster) against all allele clusters present in that sample and computed pairwise distances. For each feature, we recorded the minimum distance to any allele cluster present in the sample. This generated a 273-dimensional vector for each sample, representing its distance to each of the 273 reference features. Stacking these vectors across all samples yielded a sample-by-feature distance matrix reflecting global TVS similarity between samples.

To quantify similarity between a sample and a unique feature, we used the Jensen–Shannon distance (*58*) between TVS ratio profiles, as this metric is more sensitive to subtle fluctuations in low-abundance TVS positions compared to correlation.

Dimensionality reduction of the resulting matrix was performed using UMAP (*59*), which better preserves local neighborhood structures and allows related individuals to cluster together while separating unrelated individuals. The top two UMAP components were visualized to illustrate the family stratification based on global TVS similarity.

### Identification of recombination and gene conversion events in family 2

Analysis of TVS patterns in Family 2 revealed clear de novo TVS blocks on both Chr. 11p and Chr. 17q in the mother, which appear to be derived from the grandmother (Figures S4B and S4D). In both cases, these new TVS blocks are positioned distal to the pre-existing TVS regions—i.e., closer to the chromosome end. Based on their location and sequence composition, we hypothesized that these blocks arose via recombination or gene conversion. To investigate this, we systematically surveyed all telomere allele groups from the grandparents, identifying candidate segments with both matching sequence and length.

For Chr. 11p, the newly acquired TVS block comprises ∼220–240bp of TTGGGG repeats. A sequence-matched segment was identified only on the grandmother’s Chr. 12p allele, which was inherited by the mother. This pattern is consistent with a gene conversion event between Chr. 11p and Chr. 12p, occurring either during meiosis or through somatic replication (Figure S7).

For Chr. 17q, the new TVS block comprises ∼170–180bp of CTAGGG repeats. A matching segment was found only on one of the grandfather’s telomere alleles—however, this allele was not inherited by the mother. This suggests a more complex recombination history, possibly involving an initial crossover event during spermatogenesis, followed by a second recombination (meiotic or mitotic) after fertilization. As a result, the mother acquired a grandfather-derived TVS block on the grandmother-transmitted Chr. 17q, despite not inheriting the original allele carrying that segment (Figure S6).

## References

1. E. H. Blackburn, Telomere states and cell fates. Nature 408, 53–56 (2000).

2. T. de Lange, How shelterin solves the telomere end-protection problem. Cold Spring Harb Symp Quant Biol 75, 167–177 (2010).

3. V. L. Makarov, Y. Hirose, J. P. Langmore, Long G tails at both ends of human chromosomes suggest a C strand degradation mechanism for telomere shortening. Cell 88, 657–666 (1997).

4. W. E. Wright, V. M. Tesmer, K. E. Huffman, S. D. Levene, J. W. Shay, Normal human chromosomes have long G-rich telomeric overhangs at one end. Genes Dev 11, 2801–2809 (1997).

5. J. D. Griffith et al., Mammalian telomeres end in a large duplex loop [see comments]. Cell 97, 503–514 (1999).

6. T. de Lange, Shelterin: the protein complex that shapes and safeguards human telomeres. Genes Dev 19, 2100–2110 (2005).

7. H. Xin et al., TPP1 is a homologue of ciliate TEBP-beta and interacts with POT1 to recruit telomerase. Nature 445, 559–562 (2007).

8. E. Abreu et al., TIN2-tethered TPP1 recruits human telomerase to telomeres in vivo. Mol Cell Biol 30, 2971–2982 (2010).

9. M. F. Kendellen, K. S. Barrientos, C. M. Counter, POT1 association with TRF2 regulates telomere length. Mol Cell Biol 29, 5611–5619 (2009).

10. S. Smith, I. Giriat, A. Schmitt, T. de Lange, Tankyrase, a poly(ADP-ribose) polymerase at human telomeres [see comments]. Science 282, 1484–1487 (1998).

11. D. Broccoli, A. Smogorzewska, L. Chong, T. de Lange, Human telomeres contain two distinct Myb-related proteins, TRF1 and TRF2. Nat Genet 17, 231–235 (1997).

12. P. Konig, L. Fairall, D. Rhodes, Sequence-specific DNA recognition by the myb-like domain of the human telomere binding protein TRF1: a model for the protein-DNA complex. Nucleic Acids Res 26, 1731–1740 (1998).

13. W. Palm, T. de Lange, How shelterin protects mammalian telomeres. Annu Rev Genet 42, 301–334 (2008).

14. S. Marcand, E. Gilson, D. Shore, A protein-counting mechanism for telomere length regulation in yeast. Science 275, 986–990 (1997).

15. B. van Steensel, T. de Lange, Control of telomere length by the human telomeric protein TRF1. Nature 385, 740–743 (1997).

16. C. W. Greider, Regulating telomere length from the inside out: The replication fork model. bioRxiv, (2016).

17. K. Ancelin et al., Targeting assay to study the cis functions of human telomeric proteins: evidence for inhibition of telomerase by TRF1 and for activation of telomere degradation by TRF2. Mol Cell Biol 22, 3474–3487 (2002).

18. D. Loayza, T. De Lange, POT1 as a terminal transducer of TRF1 telomere length control. Nature 423, 1013–1018 (2003).

19. J. Feng et al., The RNA component of human telomerase. Science 269, 1236–1241. (1995).

20. T. M. Nakamura et al., Telomerase catalytic subunit homologs from fission yeast and human. Science 277, 955–959. (1997).

21. N. W. Kim et al., Specific association of human telomerase activity with immortal cells and cancer [see comments]. Science 266, 2011–2015 (1994).

22. K. Masutomi et al., Telomerase maintains telomere structure in normal human cells. Cell 114, 241–253. (2003).

23. W. E. Wright, M. A. Piatyszek, W. E. Rainey, W. Byrd, J. W. Shay, Telomerase activity in human germline and embryonic tissues and cells. Dev Genet 18, 173–179. (1996).

24. M. Z. Levy, R. C. Allsopp, A. B. Futcher, C. W. Greider, C. B. Harley, Telomere end-replication problem and cell aging. J Mol Biol 225, 951–960. (1992).

25. Z. Kaul, A. J. Cesare, L. I. Huschtscha, A. A. Neumann, R. R. Reddel, Five dysfunctional telomeres predict onset of senescence in human cells. EMBO Rep 13, 52–59 (2012).

26. M. Armanios et al., Short telomeres are sufficient to cause the degenerative defects associated with aging. Am J Hum Genet 85, 823–832 (2009).

27. Telomeres. L. V., E. H. Blackburn, T. de Lange, Eds., (Cold Spring Harbor Laboratory Press, Plainview, 2006), pp. 576.

28. C. B. Harley, A. B. Futcher, C. W. Greider, Telomeres shorten during ageing of human fibroblasts. Nature 345, 458–460 (1990).

29. I. Flores et al., The longest telomeres: a general signature of adult stem cell compartments. Genes Dev 22, 654–667 (2008).

30. I. Flores, R. Benetti, M. A. Blasco, Telomerase regulation and stem cell behaviour. Curr Opin Cell Biol 18, 254–260 (2006).

31. K. Collins, J. R. Mitchell, Telomerase in the human organism. Oncogene 21, 564–579 (2002).

32. B. Holohan, W. E. Wright, J. W. Shay, Cell biology of disease: Telomeropathies: an emerging spectrum disorder. J Cell Biol 205, 289–299 (2014).

33. M. Armanios, E. H. Blackburn, The telomere syndromes. Nat Rev Genet 13, 693–704 (2012).

34. J. Graakjaer et al., Allele-specific relative telomere lengths are inherited. Hum Genet 119, 344–350 (2006).

35. D. M. Baird, J. Rowson, D. Wynford-Thomas, D. Kipling, Extensive allelic variation and ultrashort telomeres in senescent human cells. Nat Genet 33, 203–207 (2003).

36. K. Karimian et al., Human telomere length is chromosome end-specific and conserved across individuals. Science 384, 533–539 (2024).

37. S. E. Sanchez et al., Digital telomere measurement by long-read sequencing distinguishes healthy aging from disease. Nature communications 15, 5148 (2024).

38. T. T. Schmidt et al., High resolution long-read telomere sequencing reveals dynamic mechanisms in aging and cancer. Nature communications 15, 5149 (2024).

39. C. Y. Tham et al., High-throughput telomere length measurement at nucleotide resolution using the PacBio high fidelity sequencing platform. Nature communications 14, 281 (2023).

40. Z. Stephens, J. P. Kocher, Characterization of telomere variant repeats using long reads enables allele-specific telomere length estimation. BMC bioinformatics 25, 194 (2024).

41. L. v. d. Maaten, G. Hinton, Visualizing data using t-SNE. Journal of machine learning research 9, 2579–2605 (2008).

42. R. J. G. B. Campello, D. Moulavi, J. Sander. (Springer Berlin Heidelberg, Berlin, Heidelberg, 2013), pp. 160–172.

43. S. Nurk et al., The complete sequence of a human genome. Science 376, 44–53 (2022).

44. W. W. Liao et al., A draft human pangenome reference. Nature 617, 312–324 (2023).

45. G. Palsson et al., Complete human recombination maps. Nature 639, 700–707 (2025).

46. A. B. Stergachis, B. M. Debo, E. Haugen, L. S. Churchman, J. A. Stamatoyannopoulos, Single-molecule regulatory architectures captured by chromatin fiber sequencing. Science 368, 1449–1454 (2020).

47. A. Jha et al., DNA-m6A calling and integrated long-read epigenetic and genetic analysis with fibertools. Genome Res 34, 1976–1986 (2024).

48. N. Altemose et al., DiMeLo-seq: a long-read, single-molecule method for mapping protein-DNA interactions genome wide. Nat Methods 19, 711–723 (2022).

49. H. S. Bender et al., Extreme telomere length dimorphism in the Tasmanian devil and related marsupials suggests parental control of telomere length. PLoS One 7, e46195 (2012).

50. D. Porubsky et al., Human de novo mutation rates from a four-generation pedigree reference. Nature 643, 427–436 (2025).

51. H. J. Jeon, M. T. Levine, M. A. Lampson, A parent-of-origin effect on embryonic telomere elongation determines telomere length inheritance. Curr Biol, (2025).

52. M. T. Hemann, M. A. Strong, L. Y. Hao, C. W. Greider, The shortest telomere, not average telomere length, is critical for cell viability and chromosome stability. Cell 107, 67–77 (2001).

53. F. Gao et al., Exacerbation of cardiovascular ageing by diabetes mellitus and its associations with acyl-carnitines. Aging 13, 14785–14805 (2021).

54. C. C. Liu et al., Distinct Responses of Stem Cells to Telomere Uncapping-A Potential Strategy to Improve the Safety of Cell Therapy. Stem Cells, (2016).

55. W. Shen, B. Sipos, L. Zhao, SeqKit2: A Swiss army knife for sequence and alignment processing. Imeta 3, e191 (2024).

56. C. Jain et al., Weighted minimizer sampling improves long read mapping. Bioinformatics 36, i111–i118 (2020).

57. C. Jain, A. Rhie, N. F. Hansen, S. Koren, A. M. Phillippy, Long-read mapping to repetitive reference sequences using Winnowmap2. Nat Methods 19, 705–710 (2022).

58. M. A. Re, R. K. Azad, Generalization of entropy based divergence measures for symbolic sequence analysis. PLoS One 9, e93532 (2014).

59. E. Becht et al., Dimensionality reduction for visualizing single-cell data using UMAP. Nat Biotechnol, (2018).

